# Glioma-neuronal circuit remodeling induces regional immunosuppression

**DOI:** 10.1101/2023.08.04.548295

**Authors:** Takahide Nejo, Saritha Krishna, Christian Jimenez, Akane Yamamichi, Jacob S. Young, Senthilnath Lakshmanachetty, Tiffany Chen, Su Su Sabai Phyu, Hirokazu Ogino, Payal Watchmaker, David Diebold, Abrar Choudhury, Andy G. S. Daniel, David R. Raleigh, Shawn L. Hervey-Jumper, Hideho Okada

**Affiliations:** Department of Neurological Surgery, University of California, San Francisco, San Francisco, CA, 94158, United States; Department of Radiation Oncology, University of California, San Francisco, San Francisco, CA, 94158, United States; Department of Pathology, University of California, San Francisco, San Francisco, CA, 94158, United States; Weill Institute for Neurosciences, San Francisco, CA, 94158, United States; Parker Institute for Cancer Immunotherapy, San Francisco, CA, 94129, United States

## Abstract

Neuronal activity-driven mechanisms impact glioblastoma cell proliferation and invasion^1–7^, and glioblastoma remodels neuronal circuits^8,9^. Distinct intratumoral regions maintain functional connectivity via a subpopulation of malignant cells that mediate tumor-intrinsic neuronal connectivity and synaptogenesis through their transcriptional programs^8^. However, the effects of tumor-intrinsic neuronal activity on other cells, such as immune cells, remain unknown. Here we show that regions within glioblastomas with elevated connectivity are characterized by regional immunosuppression. This was accompanied by different cell compositions and inflammatory status of tumor-associated macrophages (TAMs) in the tumor microenvironment. In preclinical intracerebral syngeneic glioblastoma models, CRISPR/Cas9 gene knockout of Thrombospondin-1 (TSP-1/*Thbs1*), a synaptogenic factor critical for glioma-induced neuronal circuit remodeling, in glioblastoma cells suppressed synaptogenesis and glutamatergic neuronal hyperexcitability, while simultaneously restoring antigen-presentation and pro-inflammatory responses. Moreover, TSP-1 knockout prolonged survival of immunocompetent mice harboring intracerebral syngeneic glioblastoma, but not of immunocompromised mice, and promoted infiltrations of pro-inflammatory TAMs and CD8+ T-cells in the tumor microenvironment. Notably, pharmacological inhibition of glutamatergic excitatory signals redirected tumor-associated macrophages toward a less immunosuppressive phenotype, resulting in prolonged survival. Altogether, our results demonstrate previously unrecognized immunosuppression mechanisms resulting from glioma-neuronal circuit remodeling and suggest future strategies targeting glioma-neuron-immune crosstalk may open up new avenues for immunotherapy.

---

Despite advances in the surgical and medical treatments for glioblastoma, the most common and aggressive malignant primary brain neoplasm, patients still face dismal prognoses^10^. Recent advancements have shed light on a previously unrecognized mechanism whereby neuronal activity drives glioblastoma growth^1–5^ and invasion^6,7,9^, through direct synaptic connections between neurons and glioblastoma cells^3,6^ as well as paracrine growth factors from glioblastoma cells and excitatory neurons^1,2,4,5^. Conversely, glioblastoma cells induce neuronal hyperexcitability and neuronal circuit hypersynchrony^11–14^.

The amount of functional connectivity between glioblastoma cells and the normal brain circuits negatively impacts patient survival through the tumor-derived synaptogenic factor thrombospondin-1 (TSP-1, encoded by the *Thbs1* gene)^8,15^. Furthermore, patient-derived glioblastoma cells from functionally connected intratumoral regions are characterized by a proliferative and invasive phenotype in the presence of neurons. The fundamental discovery that tumor microenvironment regulates malignant growth in an activity-dependent manner raises questions about additional cellular factors that may also alter or drive glioblastoma proliferation. Intriguingly, single-cell RNA-sequencing analysis on patient tumor samples has revealed that TSP-1 is predominantly expressed by glioblastoma cells within highly functionally connected (termed HFC) intratumoral regions, while non-tumor cells, including astrocytes and myeloid cells, emerge as the primary sources of the TSP-1 expression in the lowly functionally connected (termed LFC) regions^8^. This observation suggests distinct gene expression programs between HFC and LFC intratumoral regions. Myeloid cells, represented by microglia, emerge in early development, respond to the local environment by altering their molecular and phenotypic states, and regulate neuronal activity^16,17^. Moreover, glioblastoma cells and tumor-associated macrophages (TAMs) engage in bidirectional crosstalk, where glioblastoma cells attract TAMs and the TAMs promote glioblastoma cell proliferation and invasion^18,19^, preventing T-cell-mediated immune attack^20^. It is therefore possible that activity-dependent glioblastoma proliferation is governed by crosstalk between neurons, immune cells, and glioblastoma cells, which remains poorly understood.

This knowledge gap is of paramount importance to address, as gaining a deeper understanding of immune-modulating mechanisms holds the potential to unlock new therapeutic opportunities. Despite extensive efforts, and unlike the clinical success observed in various other malignancies over the past decade, cancer immunotherapy has yet to demonstrate efficacy for patients with glioblastoma^21,22^. Thus, identification of immune-modulating factors unique to the central nervous system could provide valuable insights for the development of distinctive approaches and pave the way for successful immunotherapeutic interventions. In this study, we discovered the causal role of glioblastoma cell-derived TSP-1 in the suppressive immune microenvironment in relation to distinct glioma-neuronal circuit remodeling and neuronal activities. Furthermore, we demonstrate that inhibiting glutamatergic excitatory signals by an FDA approved drug redirected the immune microenvironment of intracerebral glioblastoma, specifically a less suppressive phenotype of tumor-associated macrophages, resulting in prolonged survival.

## Functionally connected intratumoral regions have distinct immunological programs

To gain a deeper understanding of the distinctive transcriptional programs between tumor and immune cells from HFC and LFC regions, we re-analyzed the previously reported single-cell RNA-seq (sc-RNA-seq) datasets of clinical samples, with which the annotations as either HFC- or LFC-derived had been made based on presurgical MEG and MRI imaging analyses^8^. We performed differential gene expression (DGE) analyses followed by gene set enrichment analyses (GSEA) (**Extended Data Fig.S1**). The unbiased testing of 50 pathways from the MSigDB Hallmark collection revealed that numerous immune-related gene signatures were among the most significantly downregulated pathways in HFC compared with LFC regions. These downregulated pathways included *inflammatory response*, *interferon-α* and *-γ responses*, and *TNFα signaling via NFκB pathway* (**Fig. 1a–c**). These findings were consistently observed across multiple cell populations—tumor, myeloid, and lymphoid cells—while not in astrocytes (**Extended Data Fig.S2**). Signature scoring analyses for each cell demonstrated that the pathways of the *inflammatory response*, *interferon-γ response*, and *TNFα signaling via NFκB* were significantly downregulated in HFC region cells compared with LFC region cells across the cell types in common (**Fig. 1d–i**). Demonstrative genes consistently expressed in poorly connected LFC regions (over HFC) included *CCL2/4*, *CD83, IL1B*, *IRF8, NFKB1,* and *STAT1* (**Extended Data Fig.S3**). These findings led us to hypothesize that glioblastoma-intrinsic functional connectivity, which promotes the malignant behavior of glioblastoma, interacts with the immune regulation programs involving multiple cell types within these regions. Therefore, we aimed to investigate evidence of the relationship between immune systems and the neuronal activity in glioblastoma.

**Fig. 1.**
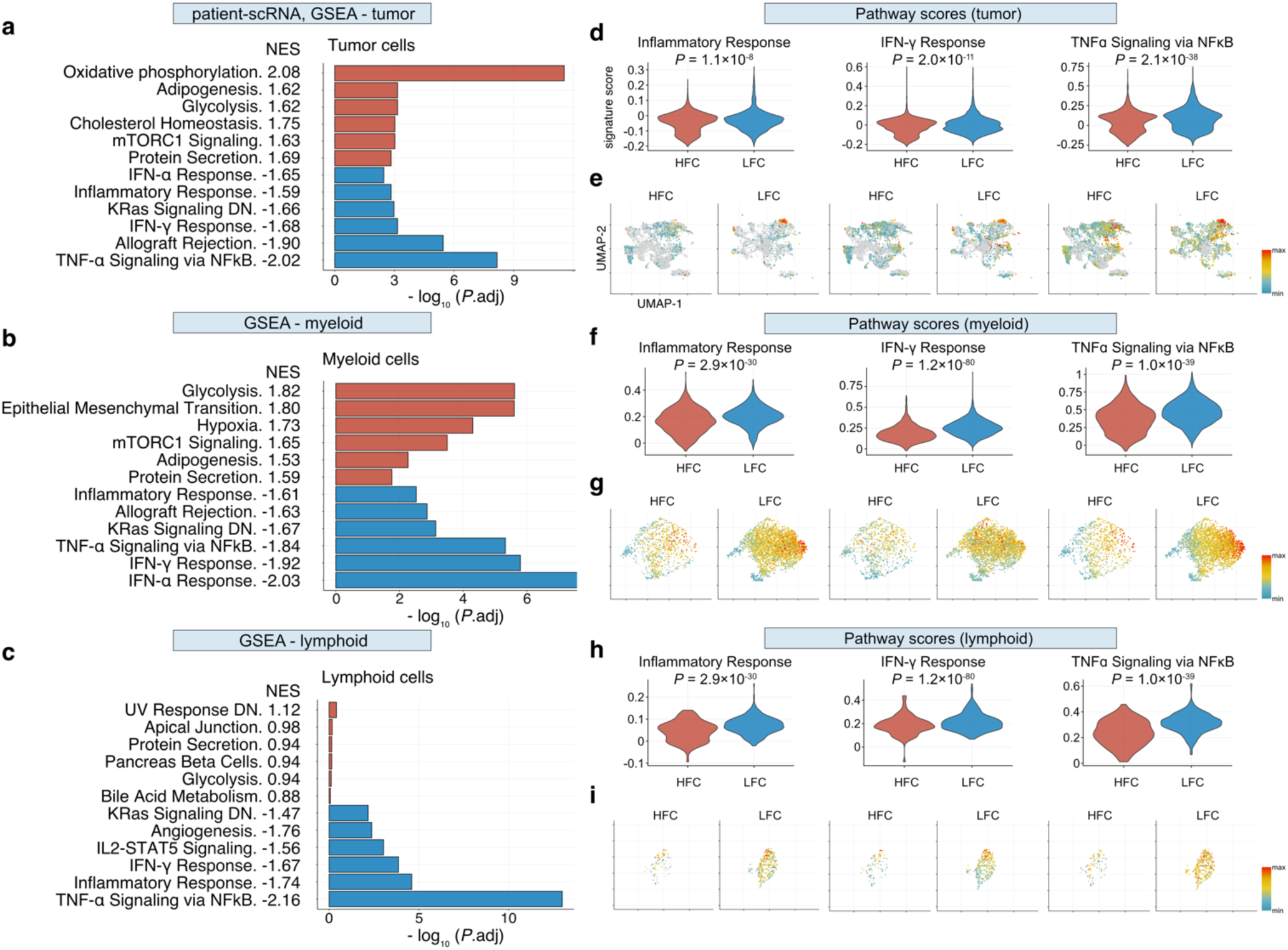
Single-cell RNA sequencing demonstrates that human glioblastoma cells derived from functionally connected intratumoral regions have distinct immune-related gene expression programs. **a–c**, Gene set enrichment analyses (GSEA) with MSigDB Hallmark collection comparing HFC vs. LFC within tumor cells (**a**), myeloid cells (**b**), and lymphoid cells (**c**). The top six upregulated and down-regulated signatures are shown for each comparison. Red and blue indicate upregulation in HFC and LFC regions, respectively. Statistical values are presented in each figure as normalized enrichment scores (NES) and adjusted *P* values. **d–i**, Violin plots (**d, f, h**), and feature plots (**e, g, i**) showing the distributions of the signature scores of *Inflammatory Response*, *Interferon-γ Response*, *TNFα signaling via NFκB pathways* (all from MSigDB Hallmark collection). *P* values are calculated using the Wilcoxon rank-sum test. In feature plots, expression values are presented as indicated in the color bar.

## Myeloid cell populations are immunosuppressive within functionally connected intratumoral regions

Myeloid cells are a major component of the glioblastoma microenvironment^23^, represented by TAMs originating from brain resident microglia (Mg-TAM) or bone-marrow monocyte-derived macrophages (Mo-TAMs)^24–26^. TAMs are typically polarized toward alternatively activated, anti-inflammatory phenotypes due to tumor-extrinsic factors, such as TGF-β, IL-10, GM-CSF, and CSF-1^27,28^, although recent studies have also indicated that TAMs in glioblastoma can exist in an immature state without showing clear polarization^19,24^. Notably, recent single-cell studies have demonstrated that, although TAMs can simultaneously express canonical classically activated (M1) and alternatively activated (M2) TAM markers on the same cells, Mo-TAMs up-regulate M2-associated immunosuppressive cytokines compared to Mg-TAMs^24^. Therefore, while their oversimplified classification remains controversial^23^, evaluating the M1 vs. M2 TAM phenotype and factors influencing the phenotype can still help us better understand the characteristics of the tumor microenvironment^24,25^. Furthermore, microglia activation regulates the activity of neurons, suggesting a potential role in glioblastoma proliferation^29^. To explore the characteristics of TAMs derived from HFC and LFC intratumoral regions, we investigated the myeloid cell populations (n = 3,775 cells) extracted from the sc-RNA-seq dataset (**Fig. 2a**).

**Fig. 2.**
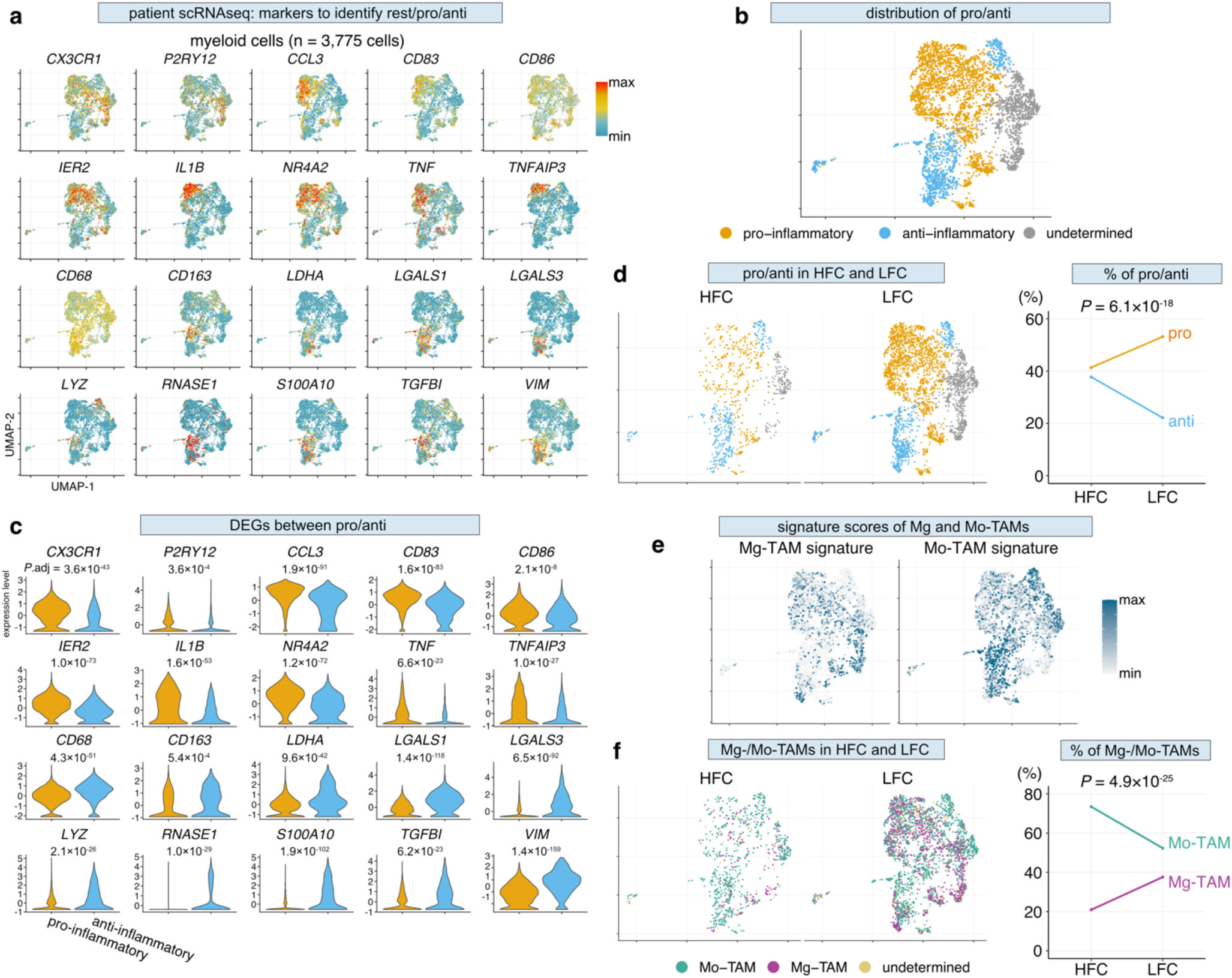
Myeloid cells derived from HFC- and LFC regions exhibit distinct activation status and populations in human glioblastoma. **a–b,** A total of 3,775 myeloid cells are analyzed. The identified myeloid subsets are annotated as either pro-inflammatory, anti-inflammatory, or undetermined, based on their distinct gene expression patterns. **c,** Violin plots confirming that representative pro- and anti-inflammatory marker genes are differentially expressed between the identified two subgroups. **d,** UMAP plots showing the relative contributions of each cell subpopulation within those isolated from intratumoral regions with HFC and LFC (**left**), and the percentages of each subpopulation within each region are shown in the line plot (**right**). **e,** Feature plots highlighting the distributions of the Mg-TAM and Mo-TAM signature scores. **f,** UMAP plots showing the relative contributions of innate immunity populations (Mg-TAM or Mo-TAM) within those isolated from intratumoral regions with HFC and LFC (**left**), and their percentages within each region are shown in the line plot (**right**). *P* values are calculated using MAST algorithm with Benjamini-Hochberg adjustment (**c**) and Fisher’s exact test (**d** and **f**).

Unsupervised clustering identified several distinct myeloid cell subclusters, including those of pro-inflammatory and anti-inflammatory status (**Fig. 2b**). Pro-inflammatory populations were represented by *CCL3*, *CD83*, *IL1B*, and *TNF*, while anti-inflammatory populations were represented by *CD163*, *LGALS1/3*, *LYZ*, *RNASE1*, *TGFBI*, and *VIM* (**Fig. 2c**). In this analysis, myeloid cells from the HFC regions demonstrated greater anti-inflammatory populations (37.7% vs. 22.1% in LFC) and less pro-inflammatory populations (41.4% vs 53.1% in LFC) compared with their LFC region cells (Fisher’s exact test *P* = 6.1×10^−18^) (**Fig. 2d**). In addition to inflammatory status, we sought to discriminate between Mg- and Mo-TAMs, given the prior observation that the compositions of innate immunity populations dynamically change along with disease progression and aggressiveness^30,31^. Signature scoring analysis demonstrated that the distributions of Mg-TAMs and Mo-TAMs overlapped with those of pro-inflammatory and anti-inflammatory TAM populations (**Fig. 2e**). As such, myeloid cells from the HFC regions were estimated to have greater Mo-TAM populations (73.4% vs. 52.3% in LFC) and less Mg-TAM populations (20.9% vs 37.6% in LFC) compared with their LFC region cells (Fisher’s exact test *P* = 4.9×10^−25^) (**Fig. 2f**). Importantly, during the presurgical clinical imaging tests used for identifying HFC and LFC regions, the HFC and LFC voxels were equally identified within both enhancing intratumoral regions as well as FLAIR hyperintense regions, and the tissues were collected accordingly^8^. Therefore, any differences observed in TAMs are unlikely to be owing to the sampling bias. Taken together, alongside the recent discovery that tumor-intrinsic neuronal activity drives glioblastoma proliferation, the significant immunosuppression within HFC intratumoral regions could be attributed to neuronal activity-mediated modulation of TAMs and their inflammatory states, where anti-inflammatory Mo-TAMs are significantly enriched.

## Inverse spatial relationship between synaptic programs and pro-inflammatory responses

Given the finding that tumor-intrinsic functional connectivity in patients with glioblastoma is heterogenous and associated with regional immunosuppression, we aimed to investigate the spatial significance of this finding *in situ*. We performed spatially-resolved transcriptomic RNA sequencing using the 10X Genomics Visium platform in a murine syngeneic glioblastoma model, SB28, which mirrors the immunological signature of human disease^32–34^. We obtained datasets consisting of 3,757 total spots (experiment performed in biological duplicates [n = 2 mice] including 1,975 and 1,782 spots, respectively) and reverse-mapped the sequencing data onto the corresponding H&E-stained 2D tissue images (**Fig. 3a** and **Extended Data Fig.S4a**). Tumor outlines were delineated based on morphology on H&E staining images as well as gene expression score distributions of *cell-cycle* (Reactome), *hypoxia* (Hallmark), and *Verhaak_Glioblastoma_Mesenchymal* (“GBM-MES” [C2:CGP]). Next, we modeled human disease by defining regions of neuronal enrichment (“glioblastoma-neuronal infiltration area”) based on the distribution patterns of the *GBM-MES* and *Neuronal Systems* (Reactome) gene enrichment scores (**Fig. 3b–e** and **Extended Data Fig.S4b–e**).

**Fig. 3.**
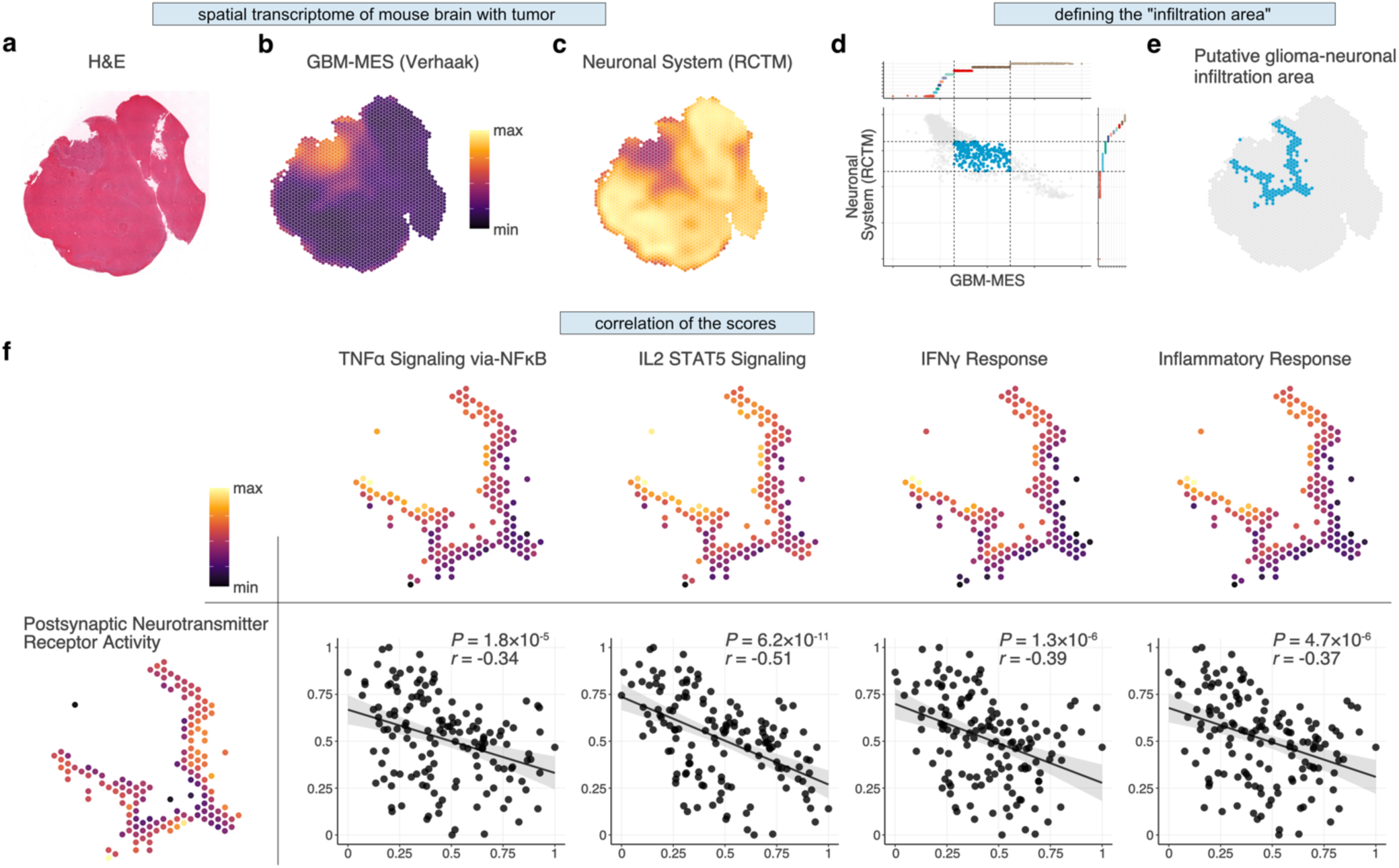
Spatial transcriptomic analyses reveal the inverse association between neuro-synaptic activities and immune regulation. **a–c,** Histological images (H&E) (**a**), and surface plots showing the distribution of the gene set enrichment scores of *Verhaak Glioblastoma Mesenchymal* (“GBM-MES” [C2:CGP]) (**b**) and *Neuronal System* (Reactome) (**c**). **d,** Scatter plot showing the relationship of the scores of *GBM-MES* and *Neuronal System* throughout the entire data set, where the spots with upper 10–30 percentiles of *GBM-ME*S scores and lower 10–30 percentiles of *Neuronal Systems* (Reactome) scores are highlighted with blue. **e,** Surface plot showing the “putative glioma-neuronal infiltration areas” defined based on the distribution of the scores of *GBM-MES* and *Neuronal Systems* within the tumor bed. **f,** Surface plots show the gene set enrichment signature scores of *Postsynaptic Neurotransmitter Receptor Activity* (GO:MF), *TNFα-Signaling via NFκB*, *IL2-STAT5 Signaling*, *IFNγ Response*, and *Inflammatory Response* (all from Hallmark) within the defined putative glioma-neuronal infiltration area. Scatter plots show the correlations between the scores of *Post-synaptic Neurotransmitter Receptor Activity* and the others. *P* values are calculated using the Pearson correlation test. *r*, correlation coefficient.

Within cortical regions with glioblastoma infiltration, we observed a strong association between synaptic related genes (represented by the gene set *Postsynaptic Neurotransmitter Receptor Activity* [GO:MF]) and downregulation of pro-inflammatory signatures (represented by the gene sets *TNF-α-via-NFκB Signaling Pathway*, *IL2-STAT5-Signaling*, *IFNγ Response*, and *Inflammatory Response* [all from Hallmark]) (**Fig. 3f**). These findings exhibited high reproducibility across the tested two individual mice (**Extended Data Fig.S4f**). In addition, deconvolution analysis using xCell validated these findings (**Extended Data Fig.S5**). Together, the spatial transcriptomic analyses provide evidence of synapse-associated gene expression within functionally connected and immunosuppressed intratumor regions across mouse and human glioblastoma models.

## TSP-1 KO reprograms tumor microenvironment to be less immunosuppressive

Recent evidence identified TSP-1 as a driver of glioma-neuron interactions within HFC regions of glioblastoma^8^. This synaptogenic factor is expressed by astrocytes in intratumoral regions without neuronal activity (LFC regions), whereas malignant tumor cells are its primary source in HFC regions. TSP-1 is multi-functional, suggesting potential roles in synaptogenesis, neuronal development^35,36^, and immunomodulation in both health and disease states^37–39^. Therefore, we hypothesized a potential causal link between glioblastoma synaptic enrichment and co-occurring immunosuppression through the production of synaptogenesis-associated paracrine-mediated factors.

To experimentally interrogate neuronal-activity-associated immunosuppression in glioblastoma, we aimed to establish a syngeneic glioblastoma model recapitulating HFC and LFC glioblastoma. We screened the three publicly available murine RNA-sequencing data (*in vitro* SB28 and GL261 syngeneic glioma cell lines and bulk normal mouse brain) and found that endogenous gene expression of TSP-1 (*Thbs1*) was significantly higher in SB28 compared with the other two datasets (vs. GL261, log_2_(fold change [FC]) = 4.50, adjusted *P* = 7.4×10^−60^; vs. normal mouse brain, log_2_(FC) = 9.54, adjusted *P* < 1×10^−300^) (**Fig. 4a**). Therefore, we hypothesized that SB28 tumors could be HFC-like because of its high expression of TSP-1, and that its downregulation could redirect the tumor characteristics toward LFC-like behavior. We generated SB28-TSP-1-knock-out (KO) clones using CRISPR-Cas9 (**Extended Data Fig.S6**), followed by single-cell cloning. The clone that underwent nucleofection without sgRNAs was used as the wildtype control (WT, “Cas9 only”). Their KO status was confirmed at the genomic DNA, mRNA, as well as protein levels (**Extended Data Fig.S7**, **S8, and S9a**).

**Fig. 4.**
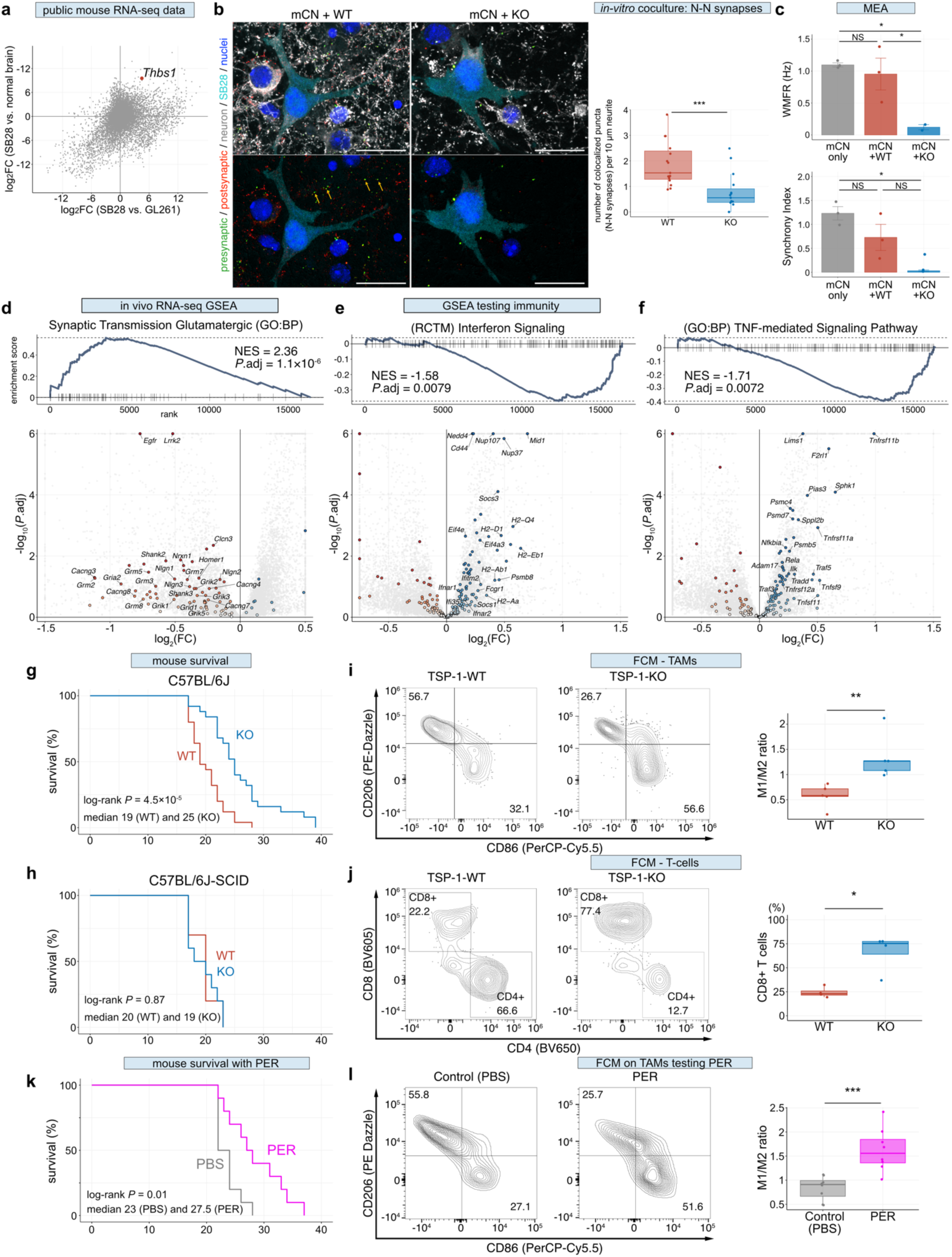
TSP-1-mediated regional immunosuppression in glioblastoma HFC regions are reversible by treatment targeting glutamatergic excitatory signals. **a,** Scatter plot illustrating the differential gene expression patterns in three separate RNA sequencing datasets: SB28 (GSE127075), GL261 (GSE94239), and normal murine brain (E-MTAB-6081). The X- and Y-axes depict log_2_ Fold Change values calculated using DESeq2. The *Thbs1* gene is highlighted in red. **b**, Representative confocal images of mouse neonatal cortical neurons co-cultured with SB28-TSP-1-WT or KO tumor cells for 24 hours showing regions of synaptic puncta colocalization (arrows in bottom panels). Red, synapsin-1 (presynaptic puncta); green, Homer-1 (post-synaptic puncta); white, Tubulin-β III (TUJ, neurons); cyan, GFP (SB28 tumor cells); blue, DAPI. Scale bar, 20 µm. Box plot showing quantification of the number of colocalized pre- and post-synaptic puncta identified on neurites (n = 15 ROIs per group, *P* = 7.8×10^−4^. Median: 1.94 [WT] vs. 0.62 [KO] per 10 µm neurite). **c,** Bar plots showing the MEA data comparing mouse neonatal cortical neurons alone (mCN only), neurons co-cultured with SB28-TSP-1-WT or KO tumor cells (mCN + WT and mCN + KO, respectively). Neuronal activity amplitudes and the synchronies are quantified and represented as the weighted mean firing rate (WMFR)(Hz) and network synchrony (synchrony index) measured at 24 hours post co-culture. WMFR: *P* = 0.55 (mCN only vs. mCN + WT); *P* = 0.03 (mCN only vs. mCN + KO); *P* = 0.03 (mCN only vs. mCN + KO). Synchrony index: *P* = 0.13 (mCN only vs. mCN + WT); *P* = 0.03 (mCN only vs. mCN + KO); *P* = 0.11 (mCN only vs. mCN + KO). Data is acquired through technical replicates as follows: n = 3 wells for mCN only and mCN + WT co-culture; n = 2 wells for mCN + KO co-culture. Data are mean ± s.e.m. **d–f**, Enrichment plots and volcano plots summarizing GSEA with the gene sets *Synaptic Transmission Glutamatergic* (GO:BP) (**d**), *Interferon Signaling* (Reactome) (**e**), and *TNF-mediated Signaling Pathway* (GO:BP) (**f**), to compare the gene expression patterns between the SB28-TSP-1-WT and KO mouse tumors. In enrichment plots, positive and negative normalized enrichment scores (NES) indicate the upregulation and downregulation in WT tumors compared to the KO counterparts, respectively. In volcano plots, the genes composing the gene set are highlighted in colors, and among them, the representative leading-edge gene symbols are labeled. Genes with log_2_FC values and adjusted *P* values exceeding the boundaries are flattened and shown on the edges. **g–h,** Kaplan-Meier survival curves for the SB28-TSP-1-WT or KO (10,000 cells/1 µL/mouse) inoculated into C57BL/6J immunocompetent mice (n = 25 mice each) (**g**), or immunocompromised C57BL/6J-SCID mice (n = 10 mice each) (**h**). **i–j,** Contour plots and box plots summarizing the flow cytometry data performed on the brain-infiltrating leukocytes isolated from the mouse brain harboring SB28-TSP-1-WT or KO tumors on day 20 after tumor cell inoculation. **i,** Contour plots depict the distribution of “classically activated” pro-inflammatory (CD86+CD206-) and “alternatively activated” anti-inflammatory (CD86-CD206+) populations of CD45+CD11b+F4/80+ tumor-associated macrophages/microglia (TAMs). The corresponding box plot shows the ratio of CD86+/CD206- to CD86-/CD206+ (termed M1-to-M2 ratio) calculated based on the percentages of each subpopulation identified in each sample (n = 5 mice per group. *P* = 0.008. Median, 0.58 [WT] vs. 1.27 [KO]). **j,** Contour plots of representative cases from each group distinguishing CD4-CD8+ and CD4+CD8-populations within CD45+CD3+ T-cells. The corresponding box plot shows the percentages of CD8+ T-cells identified in each sample (n = 4 mice per group. *P* = 0.03. Median, 23.0% [WT] vs. 75.3% [KO]). Values in the plots indicate the percentages of the cell populations identified within the gate. **k,** Kaplan-Meier survival curves of the C57BL/6J immunocompetent mice orthotopically inoculated with SB28-TSP-1-WT cells (10,000 cells/1 µL/mouse), and treated with perampanel (PER), or vehicle control (PBS) (n = 10 mice per group) from the next day of tumor inoculation. **l,** Contour plots and box plots summarizing the flow cytometry data performed on the brain-infiltrating leukocytes isolated from the tumor-harboring mouse brains on day 20 after tumor cell inoculation. The mice are treated with PER or PBS from the next day of tumor inoculation until the euthanasia. Contour plots depict the distribution of “classically activated” pro-inflammatory (CD86+CD206-) and “alternatively activated” anti-inflammatory (CD86-CD206+) populations of CD45+CD11b+F4/80+ tumor-associated macrophages/microglia (TAMs). The corresponding box plot shows the M1-to-M2 ratio calculated based on the percentages of each subpopulation identified in each sample (n = 8 mice per group. *P* = 6.2×10^−4^. Median, 0.91 [WT] vs. 1.56 [KO]). *P* values are calculated using Wilcoxon rank-sum test (**b, i–j, l**), two-tailed Student’s t-test and adjusted using Benjamini-Hochberg method (**c**), and log-rank test (**g, h, k**). \**P* < 0.05, \*\**P* < 0.01, \*\*\**P* < 0.001; NS, not significant. Box-and-whisker plots display the median (center line) and interquartile range [IQR] (box limits). The upper whisker extends to the minimum of either the maximum value or upper quartile + 1.5×IQR, while the lower whisker extends to the maximum of either the minimum value or lower quartile - 1.5×IQR.

Next, to characterize the impact of TSP-1 on bidirectional interaction between SB28 glioblastoma cells and mouse cortical neurons *in vitro*, we analyzed immunofluorescence staining using confocal microscopy (**Fig. 4b**) ^8^. In these samples, tubulin-βIII marks neurons, synapsin-1 marks presynaptic puncta, homer-1 marks postsynaptic puncta, and SB28 glioblastoma cells are labeled by GFP. We found that the number of co-localized presynaptic and postsynaptic puncta significantly decreased in the co-culture with SB28-TSP-1-KO cells compared with those with the WT (*P* = 1.3×10^−6^, median 1.94 [WT] vs. 0.62 [KO] per 10 µm neurite). Moreover, the electrophysiological properties of SB28-TSP-1-WT and KO cells were analyzed in co-culture with neurons using a multi-electrode array (MEA). After 24 h of co-culture, the total number of network bursts (a measure of neuronal activity) and network synchrony remained unchanged in the co-culture with SB28-TSP-1-WT compared with the condition of cortical neuron only (baseline). In contrast, the KO cells showed significant decreases in both the network bursts (vs. baseline, *P* = 0.03; vs. WT, *P* = 0.03) and the synchrony (vs. baseline, *P* = 0.03), ascertaining the critical role of TSP-1 in neuronal network activity (**Fig. 4c** and **Extended Data Fig.S10**) ^8^.

We further characterized the SB28-TSP-1-WT and KO tumors *in vivo* as syngeneic orthotopic models and their interactions with surrounding non-tumor cells, including neurons. Quantitative analysis of immunofluorescence tissue staining confirmed that TSP-1 protein expression was significantly downregulated in the KO tumors (*P* = 0.008) (**Extended Data Fig.S11**). Next, we performed bulk RNA-seq on the resected tumor tissues (**Extended Data Fig.S9b–c**, **S12**). Importantly, DGE analyses followed by GSEA revealed that TSP-1-WT tumors were significantly enriched for genes associated with synapse and circuit assembly compared to the TSP-1-KO counterparts, as represented by *Neuronal System* (Reactome) and *Synaptic Transmission Glutamatergic* (GO:BP) (**Fig. 4d** and **Extended Data Fig.S13a–b**). Indeed, among the ~6,300 gene sets we tested, the top gene sets enriched in WT tumors were occupied by those related to neural activity and synapses, emphasizing the importance of TSP-1 in neuro-synaptic activities. These findings collectively demonstrated that SB28-TSP-1-WT and KO models recapitulated critical characteristics of human glioblastoma HFC and LFC intratumoral regions.

Notably, GSEA also revealed the significant restoration of immune-related signatures in the KO tumors, such as *interferon signaling*, *antigen processing and presentation via MHC-class-I*, and *TNF-mediated signaling pathways*, compared with the WT counterparts (**Fig. 4e–f** and **Extended Data Fig.S13c**). Representative DGEs upregulated in the KO tumors included *Cd44, Tnfrsf11a/b, Nfkbia, Tap1/2,* and numerous MHC class-I/II genes, interferon-related genes, and proteasome-related genes.

We also investigated the impact of TSP-1 knockout on *in vivo* tumor growth and overall survival of the mice. First, we performed stereotactic inoculation of SB28-TSP-1-WT or KO cells into immunocompetent C57BL/6J mice and found that the mice bearing the KO tumor showed significantly slower tumor growth and prolonged survival compared with their WT counterparts (19 days [WT] vs. 25 days [KO], log-rank test *P* = 4.5×10^−5^) (**Fig. 4g** and **Extended Data Fig.S14a**). Next, to investigate the contributions of adaptive immunity to the observed survival differences, we repeated the same experiment with C57BL/6J-derived severe combined immune deficient (SCID) mice that lack T-cell and B-cell systems. Notably, the differences in the tumor growth and survival were completely abrogated (20 days [WT] vs. 19 days [KO], log-rank test *P* = 0.87) (**Fig. 4h** and **Extended Data Fig.S14b**). These findings collectively indicate that the survival difference observed in the immunocompetent animals is attributable to the role of adaptive immunity.

Therefore, we next investigated brain-infiltrating leukocytes (BILs) isolated from tumor-harboring mouse hemispheres, using flow cytometry with 1) myeloid- and 2) T-cell-markers. We defined CD45+CD11b+F4/80+ cells as tumor-associated macrophages/microglia (TAMs) and characterized them by staining with CD86 and CD206. In contrast to TAMs isolated from the WT tumors, which predominantly exhibited the CD86-CD206+ “alternatively activated” phenotype, the cells isolated from the KO tumors consistently demonstrated polarization towards the CD86+CD206-“classically activated” phenotype (n = 5 mice in each group) (**Fig. 4i**, and **Extended Data Fig.S15**). Correspondingly, the ratio of CD86+/CD206- to CD86-/CD206+ (termed M1-to-M2 ratio) was significantly higher in the TAMs isolated from the KO tumors compared with their WT counterparts (median: 0.59 vs. 1.34; *P* = 0.008), indicating a shift in the tumor microenvironment towards less immunosuppressive and more pro-inflammatory as a result of downregulation of TSP-1.

Furthermore, we also characterized the CD45+CD3+ tumor-infiltrating T-cell populations. The limited number of cells isolated may have impeded the comprehensive characterization of T-cells in the SB28 tumors, as these tumors are intrinsically characterized by sparse T-cell infiltration, rendering them immunologically cold^33,34^. Nevertheless, the analysis unveiled a marked disparity in the cellular composition between the WT and KO tumors. Of note, the CD3+ T-cells isolated from the KO tumors exhibited significantly higher abundances (median, 95.5 vs. 516; *P* = 0.03) and percentages (median: 23.0% vs. 75.3%; *P* = 0.03) of CD8+ T-cells within the total CD3+ T-cell population compared to WT counterparts (**Fig. 4j** and **Extended Data Fig.S16**, **S17**). In both groups, over 95% of the CD3+, CD4+, and CD8+ T-cell populations displayed an effector or effector-memory phenotype (**Extended Data Fig.S18a–c**). No difference was detected regarding the fraction of CD25+ cells within the CD4+T-cell populations between the two groups, as well as the tested activation and exhaustion markers, such as PD-1 (**Extended Data Fig.S18d**). These findings collectively demonstrate that TSP-1 knockout not only suppresses glioma-associated hyperexcitability but also restores immune responses through the increase of “classically activated” pro-inflammatory TAMs and CD8+ T-cells in the tumor microenvironment of glioblastoma, ultimately leading to prolonged survival.

## Therapeutic targeting of AMPAR rescues immunosuppression

Analyses of human clinical data and subsequent investigations in mouse preclinical models, including gene perturbation studies, underscore the critical role of tumor-derived TSP-1 in glioma-associated hyperexcitability and the accompanied regional immunosuppression. Furthermore, analysis of bulk RNA-seq of *in vivo* tumors comparing the TSP-1-WT and KO tumors revealed a significant upregulation of glutamatergic signaling pathways in the WT tumors (**Fig. 4d**) as well as significantly higher AMPAR gene expression scores^40^ compared with the KO counterparts (*P* = 0.02) (**Extended Data Fig.S19a**). Although both glutamatergic and GABAergic signals were upregulated in the WT tumors, notable shifts of glutamatergic signaling pathways were more prominent than the GABAergic signal (**Extended Data Fig.S19b–e**). Therefore, to test the hypothesis that excitatory neuronal synaptic programs drive immunosuppression, we chose perampanel (PER), an FDA-approved, anti-epileptic drug currently used in clinical practice, for evaluation in an *in-vivo* syngeneic orthotopic model. We treated the C57BL/6J mice bearing SB28-TSP-1-WT tumor with PER or vehicle from day one post-inoculation until the onset of signs necessitating euthanasia. Intriguingly, treatment with PER significantly prolonged overall survival (23 days [vehicle] vs. 27.5 days [PER], log-rank test *P* = 0.01) (**Fig. 4k**). Moreover, flow cytometry analysis of the BILs harvested on day 20 showed that the CD45+CD11b+F4/80+ TAM populations isolated from the brain of PER-treated mice were less polarized toward the “alternatively activated” phenotype and more polarized toward the “classically activated” phenotype compared to the control group (M1-to-M2 ratio median: 0.91 vs. 1.56; *P* = 6.2×10^−4^) (**Fig. 4l** and **Extended Data Fig.S20**), similar to the observed difference between the TSP-1-WT and KO tumors. Collectively, these findings highlight the potential opportunities for neuronal activity-oriented therapeutic interventions, such as PER, to redirect the tumor microenvironment of glioblastoma to be less immunosuppressive.

## Discussion

In this study, we uncover previously unrecognized evidence of regional immunosuppression within the cortex remodeled by glioblastoma infiltration, including distinct polarization and cell composition patterns of TAMs. Spatial transcriptomic analysis in a preclinical glioblastoma model reveals an inverse relationship between neuronal activity-related signatures and inflammatory response signatures. Importantly, our preclinical investigations demonstrate that TSP-1 knockout in glioblastoma cells not only suppresses neuronal activity, but also restores pro-inflammatory responses and antigen-presentation machinery, promoting the infiltration of pro-inflammatory TAMs and CD8+ T-cells within the tumor microenvironment, resulting in prolonged survival. These findings underscore the significance of TSP-1 expressing synaptogenic glioblastoma cells in dictating neuronal circuit remodeling and neuro-synaptic activity, and regional immunosuppression. Furthermore, we show that pharmacological inhibition of glutamatergic excitatory signals redirects TAMs toward a less immunosuppressive state, leading to prolonged survival.

Advancements in cancer neuroscience research have revealed unique features and treatment resistance mechanisms specific to brain tumors^1–9^. Furthermore, the contributions of the immune axis to neurofibromatosis-1 (NF1) low-grade glioma growth has also been investigated^41^. In this context, the present study sheds light on the distinctive tumor immune microenvironment resulting from enhanced synaptic connections and neuronal activities during neuronal circuit remodeling in glioblastoma. Our interdisciplinary investigation incorporates cancer neuroscience, cancer immunology, and neuroimmunology to gain comprehensive insights. Neurons, particularly during neurodevelopmental stages, actively interact with and regulate other cell types, such as microglia^17,42^ and astrocytes^35,43,44^, shaping the mature neuronal microenvironment^45^. Existing literature on neuron-microglia interactions aligns with our findings, emphasizing the critical role of proper immune response regulation, including an anti-inflammatory phenotype of microglia/macrophages, for neuronal protection^46^. Malignant cells, such as glioblastoma, can exploit any available biological programs, including those associated with developmental stages. Therefore, it is highly possible that glioblastomas govern the neuronal circuits and reprogram the immune tumor microenvironment to their advantage, evading immune attack and promoting proliferation and invasion.

Despite the progress made in our study, several important questions remain unanswered. Firstly, the involvement of other cell types, in regulating inflammatory responses needs further evaluation based on recent studies^47,48^. Secondly, the specific neuronal activity-related molecules responsible for immunomodulation have yet to be identified. Our *in vivo* observations demonstrate the favorable effects of PER treatment on mouse survival and TAM polarization towards a less immunosuppressive state, leading to the hypothesis that the effect of TSP-1 on the immune tumor microenvironment in glioblastoma is not direct but indirect, through a neuronal activity-dependent manner. Further investigations are necessary to address the hypothesis and to identify the precise molecules that directly influence the recruitment and phenotype of immune cells, such as TAMs. Thirdly, to advance cancer immunotherapy against glioblastoma, we are particularly interested in exploring the potential of therapeutic interventions targeting neuronal activity to enhance the efficacy of immune cell therapies. While our data demonstrate promising reprogramming of the immunosuppressive tumor microenvironment, further examination is needed to assess the strength of its impact. Addressing these questions will deepen our understanding of the mechanisms underlying glioma-neuron-immune crosstalk and open new avenues for cancer immunotherapy in combination with strategies targeting neuronal activity in glioblastoma.

## Methods

### Study approval

For all human tissue studies, written informed consent was obtained from all patients, and tissue samples were used in accordance with the University of California, San Francisco (UCSF) institutional review board (IRB) for human research as previously described^8^. All the experiments and analyses using clinical samples were conducted according to the Declaration of Helsinki. All the mouse studies were performed following the protocol approved by the Institutional Animal Care and Use Committee (IACUC) of UCSF.

### Single-cell gene-expression data processing and analysis

Single-cell RNA sequencing on patient clinical samples was performed as previously reported. The data has been deposited at the NCBI Gene Expression Omnibus and made publicly available under the accession code GSE223063. The resulting FASTQ files were processed using CellRanger v3.0.2 (10x Genomics) for alignment to the hg38 reference genome. The resulting filtered count matrix was further processed using R (v4.1.2) and R package Seurat (v4.0.3), including normalization and scaling using SCTransform^49^, batch-effect correction using Harmony^50^, and dimensional reduction with PCA and UMAP, as previously reported^8^. Based on the previously defined cell type annotations, in the present study, the following four clusters were subsetted and analyzed separately: tumor, myeloid, and lymphoid cells, and astrocytes. After removing hemoglobin- and ribosome-related genes from the dataset, raw count data of each subset was normalized and scaled again. Differential gene expression analyses were performed within each cell type to compare HFC and LFC using a hurdle model tailored to scRNA-seq data, part of the MAST software package^51^. The output was sorted based on the stat values to prepare the rank object. Preranked gene set enrichment analysis (GSEA) was performed using fgsea (1.18.0)^52^ for the gene set collection Hallmark (msigdbr 7.4.1). To visualize the data in violin plots and feature plots, signature scores for *Inflammatory Response*, *Interferon-gamma Response*, and *TNF-alpha Signaling via NFκB* (all from the Hallmark collection) were calculated using the AddModule() function of Seurat package. All genes within each gene set were included as input features. In addition, in-depth characterization analysis was performed for the myeloid cell populations (n = 3,775 cells). Clusters were identified using shared nearest neighbor-based (SNN-based) clustering using the first 30 principal components with k = 30 and resolution = 0.2. A total of 6 clusters were initially identified, and then manually curated into 3 clusters (pro-inflammatory, anti-inflammatory, and undetermined) based on the expression of known marker genes^24,26^. Signature scores for Mg-TAM and Mo-TAM were calculated using the AddModule() function of Seurat package. The gene expression data of *P2RY12*, *CX3CR1*, *NAV3*, *SIGLEC8*, *SLC1A3* were used for Mg-TAM, and those of *TGFBI*, *ITGA4*, *IFITM2*, *FPR3*, *S100A11*, *KYNU* were used for Mo-TAM, respectively^24^. A cell was defined as a Mo-TAM or Mg-TAM when the signature score was greater than 0 and higher than the other score. If both scores were less than 0, cells were labeled as undetermined.

### Culture of tumor cell line

A C57BL/6J-background murine glioblastoma cell line SB28^32–34^ and the deriving cells (passage number 12–25) were maintained in complete RPMI [RPMI 1640 media supplemented with 10% FBS, 1% Penicillin-Streptomycin (Gibco, 15070063), 1% HEPES (Gibco, 15630080), 1% Glutamax (Gibco, 35050061), 1% MEM non-essential amino acids (Gibco, 11140076), 1% sodium pyruvate (Gibco, 11360070)]. The cell line was originally developed in our laboratory and is now distributed through the DSMZ-German Collection of Microorganisms and Cell Cultures (https://www.dsmz.de/collection/catalogue/details/culture/ACC-880). Cells were passaged when reaching subconfluent every 3–4 days using Accutase (AT104, Innovative Cell Technologies), and maintained in a humidified incubator in 5% CO2 at 37°C. All cells were routinely confirmed to be negative for mycoplasma infection every 3–4 months using PlasmoTest mycoplasma detection kit (InvivoGen, catalog # rep-pt1). No other authentication assay was performed.

### Orthotopic mouse glioblastoma models

All mouse experiments were performed following the protocol approved by IACUC of UCSF. For orthotopic syngeneic models, SB28 tumor cells were implanted intracerebrally into 5–7 week-old female C57BL/6J mice (Jackson Laboratory, 000664) or B6.Cg-*Prkdc^scid^*/SzJ (“C57BL/6J-SCID”) mice (Jackson Laboratory, 001913) with 8–10 mice per group. The surgical procedure used in the current study has been described previously^53,54^. Briefly, animals were anesthetized with 1.5–3% isoflurane and placed in a stereotactic frame (Kopf). After disinfection with betadine and ethanol and making a midline scalp incision, the injection site was located 2 mm to the right of the bregma. A burr hole was drilled at the injection site using a 25G needle. A Hamilton syringe loaded with tumor cells (approximately 1×10^4^ cells in 1 µL sterile HBSS) and equipped with a 26G needle was inserted into the brain at a depth of 3.5 mm from the skull and then slowly raised to 3.0 mm to create space for the cells. The cells were injected targeting the right striatum at a speed of 1 µL/min using an autoinjector system. After the infusion, the syringe needle was held in place for 1 min, then pulled back to a depth of 1.5 mm and held in place for another min before being withdrawn slowly to minimize backflow of the injected cell suspension. The burr hole was sealed with bone wax. Aseptic techniques were used throughout the surgical procedure. Postoperatively, animals were treated with an analgesic (meloxicam and buprenorphine) and monitored for adverse symptoms in accordance with the IACUC-approved protocol.

### Bioluminescence imaging

Tumor engraftment and progression were monitored by luminescence emission on an IVIS imaging system (Xenogen). Mice were anesthetized with isoflurane and intraperitoneally (i.p.) injected with 1.5 mg of d-luciferin (GoldBio) in a total injection volume of 100 µL. The average radiance signal was used to generate all tumor growth data.

### Mouse drug treatment studies

For all drug studies, tumor cells were inoculated into the mice following the procedure described above. Drug treatment was initiated on the next day of tumor inoculation without performing tumor size-based randomization. Instead, the experimental groups were assigned based on the body weight of each animal, ensuring a similar range of body weights within the groups. The mice received systemic administration of either PBS or perampanel (0.75 mg/kg; Adooq Biosciences, A12498; formulated in 10% DMSO, 60% PEG300, 30% water) via oral gavage daily until reaching the endpoint, unless otherwise indicated.

### Mouse survival studies

For survival studies, morbidity criteria were consistent with the predetermined IACUC-approved biological endpoint, which included severe physiological symptoms (e.g., hunching, respiratory distress), severe neurological impairments (e.g., circling, ataxia, paralysis, limping, head tilt, balance problems, and seizures), and a body weight loss of 15% or more from the initial weight. Kaplan–Meier survival analysis, using log-rank testing, was employed to assess statistical significance.

### Spatial transcriptomics data acquisition

Spatial resolved transcriptomic data was acquired for the mouse brain tissues harboring SB28 tumors (n = 2 mice) using the Visium Spatial for FFPE Gene Expression Kit, Mouse Transcriptome (10x Genomics, 1000339). Tissue Optimization and Library preparation were carried out according to the manufacturer’s protocol (10X Genomics, CG000408 Rev A). Briefly, formalin-fixed, paraffin-embedded (FFPE) tissue blocks were made from mouse brain tissue harboring tumors collected immediately post-euthanasia and cardiac perfusion with PBS. Tissues were placed in 4% Paraformaldehyde for 24-h fixation and replaced with 70% EtOH until ready for processing. Tissues were processed and embedded into FFPE blocks on the Sakura VIP 6 and Tissue Tek 5 embedder, respectively, at the UCSF Neurosurgery Brain Tumor Center (BTC) Biorepository. Quality control of tissue blocks was performed by extracting RNA from FFPE samples using the Qiagen RNeasy FFPE Kit (Qiagen, 73504), followed by the assessment on the Agilent 2100 Bioanalyzer using the RNA 6000 Pico Kit (Agilent, 5067-1513). The DV200 values were confirmed to be 67% and 70%, meeting the minimum requirement of no less than 50%. Then, 5 µm-thick sections were mounted onto each spatially barcoded capture area of the Visium Spatial Gene Expression slide. The mounted slide was dried by storing in a desiccator at RT overnight, and finally at 60°C for 2 hours. Deparaffinization was performed with xylene; then, the slide was immediately stained with hematoxylin and eosin and imaged at 60X magnification at the Gladstone Institute Histology and Light Microscopy Core. The subsequent library preparation, quality control, and sequencing steps were performed at the Gladstone Institute Genomics Core. Tissue sections were de-crosslinked at 70°C for 1 hour, then hybridized overnight with the mouse transcriptome probes. Probe ligation was followed by the release of single-strand product from the tissue and binding to the Visium slide. Probe extension then added the unique molecular identifier (UMI), spatial barcode, and partial Read 1. Following the elution of samples from the Visium slide, one microliter of each sample was subjected to 25 cycles of qPCR. The optimal amplification cycles were determined using the Cq values at the exponential phase of the amplification plot, which is roughly 25% of the peak fluorescence. These values were used in the subsequent library preparation steps, where dual indexes were added to the barcoded products. Quality control of the final libraries was completed on the Agilent 2100 Bioanalyzer using the High Sensitivity DNA Kit (Agilent, 5067-4626) to determine the average library sizes, in addition to qPCR on the Applied Biosystems QuantStudio 5 Real-Time PCR System using the Roche KAPA Library Quantification Kit (Roche, KK4824) to determine the concentration of adapter-ligated libraries. Finally, libraries were pooled and sequenced on the NextSeq 500 high output 150 cycle flow cell, paired-end 28×50bp with 10bp dual indexes, resulting in greater than 44,000 paired reads per capture spot.

### Spatial transcriptomics data processing

Space Ranger v1.3.1 (10x Genomics) was used to integrate the FASTQ sequencing files and the H&E staining image files, construct initial count matrices for each unique molecular identifier (UMI) at every location in each sample using the mouse reference (refdata-gex-mm10-2020-A), and generate output files for subsequent analyses. Downstream analysis and visualization were done using R (v4.1.2) and the R package SPATA2 (v0.1.0) (https://github.com/theMILOlab/SPATA2)55. Denoising of the data was performed using the runAutoencoderDenoising() function from the package. All subsequent analyses were performed on the denoised expression data. Gene expression signature scores were calculated using AddModuleScore() function from Seurat. To perform region-specific analyses, the tumor bed was identified morpholotically as well as by referring to the distribution of gene expression signatures, such as *HM_HYPOXIA*, *CELL_CYCLE* (C5:GO:BP), and *VERHAAK_GLIOBLASTOMA_MESENCHYMAL* (“*GBM-MES*” [C2:CGP]). Then the tumor bed region was approximately delineated with adequate margins using the createSpatialSegmentation() function of SPATA2. Next, gene signature scores for *GBM-MES* and *NEURONAL_SYSTEMS* (C2:CP:Reactome) were calculated across all spots in the entire dataset. Based on their distribution patterns, spots with scores in the upper 10–30 percentiles of *GBM-MES* and lower 10–30 percentiles of *Neuronal Systems* were identified. Finally, spots located outside the tumor bed regions were excluded, and the remaining spots were defined as “putative glioma-neuronal infiltration areas”. For the spots identified as infiltration area, the gene expression signature scores for *TNFA_SIGNALING_VIA_NFKB*, *IL2_STAT5_SIGNALING*, *INTERFERON_GAMMA_RESPONSE*, *INFLAMMATORY_RESPONSE* (all from the Hallmark collection) and *POSTSYNAPTIC_NEUROTRANSMITTER_RECEPTOR_ACTIVITY* (C5:GO:MF) were recalculated using AddModuleScore() function. Cell-type deconvolution analysis was performed on the transcriptome data of each spot within the glioma-neuron infiltration area (n = 148 spots) using xCell^56^. From the output data matrix, which contained calculated scores of 66 cell types and 3 scores for each spot, we carefully curated 22 cell types, comprising 18 immune cells and 4 central nervous systems cell types (astrocytes, endothelial cells, neurons, and pericytes). Next, Pearson’s correlation scores were calculated among the 22 cell types and 6 relevant gene signature scores, as visualized in **Extended Data Fig.S5**.

### Analysis of public transcriptome data

We obtained three publicly available bulk transcriptome datasets: GSE127075 (SB28) and GSE94239 (GL261) from Sequence Read Archive (SRA), and E-MTAB-6081 (C57BL/6 normal mouse brain) from EMBL-EBI ArrayExpress (n = 3 samples each). The fastq reads were aligned to the mouse reference genome mm10 (GRCm38.p6) using STAR (v2.7.9a), with transcriptome annotation guidance from gencode.vM25.annotation.gtf. Sorting and indexing were performed using samtools (v1.14), and gene-level expression counts were estimated using stringtie (v2.0)^57^. All subsequent computational analyses were conducted using R (v4.1.2). Differential expression (DE) analyses comparing two groups were performed using the DESeq2 R package (v1.32.0)^58^. Log_2_ fold change (FC) values from each analysis were used for data visualization.

### Gene knockout using CRISPR-Cas9 system

Gene knockout was performed as previously described with some modifications^59^. Briefly, the Gene KO kit v2 (Synthego) was used according to the manufacturer’s recommended protocol. The kit contains three multi-guide sgRNAs specifically targeting regions within exon 3 (ENSMUSE00000295002) of the murine *Thbs1* gene. To prepare ribonucleoprotein (RNP) complexes, we mixed 60 pmol of multi-guide sgRNAs (20 pmol per each of the three individual gRNAs) and 20 pmol of recombinantly produced and purified Cas9-2NLS (QB3 Macrolab at QB3-Berkeley) at a molar ratio of 3:1 for sgRNA to Cas9. The mixture was incubated for 30 minutes at RT. For the TSP-1 wildtype control (an experimental negative control), only the Cas9 protein was added without sgRNAs. Electroporation was performed using SE Cell Line 4D-Nucleofector X kit S (Lonza, V4XC-1032). SB28 parental cells at a subconfluent condition were dissociated with Accutase, washed with PBS, and 150,000 cells were added to each tube containing RNP complexes. The volume was adjusted to 20 µL with Nucleofector solution and transferred to the Nucleocuvette Vessel provided in the kit. Nucleofection was performed using the DS126 program (“MG-U87”) of the 4D-Nucleofector X Unit (Lonza). Immediately after nucleofection, cells were recovered by adding 80 µL of pre-warmed growth media to each well, and then in a humidified incubator with 5% CO2 at 37°C for 30 minutes. The cells were collected, replated on culture dish plates, and allowed to grow. On day 4, cells were detached, dissociated, and subjected to limiting dilution and clonal expansion in a 96-well plate. The knock-out status of the *Thbs1* gene in the expanded clones was determined by amplifying genomic DNA using polymerase chain reaction (PCR), followed by Sanger sequencing as well as Western blotting, as described below. Clone 1C1 was selected as a representative knock-out clone based on induced frame-shift alterations at the genomic and transcriptomic levels, as well as robust downregulation at the protein level. This clone was used for all subsequent *in vitro* and *in vivo* experiments unless otherwise specified.

### Sanger sequencing

The DNeasy Blood & Tissue Kit (Qiagen, 69506) was used to extract genomic DNA from the culture cells, according to the manufacturer’s protocols. An aliquot of genomic DNA was amplified by polymerase chain reaction (PCR) using the following oligo primers. Fwd: 5’-TAAGGATGCAGCTTCCCTCG-3’; Rev: 5’-CCGTTGGAGACCACACTGAA-3’. After the gel electrophoresis and image acquisition, the agarose gel pieces were cut out and digested using NucleoSpin® Gel and PCR Clean-Up kit (Takara, 740609), according to the manufacturer’s protocols. Sequencing was performed at Quintara Biosciences, using the following sequencing primer: 5’-TTTCCATAATTGCCATTATTGTCACGAGTT-3’. Acquired Sanger sequencing data was analyzed and visualized using ApE (v2.0.61).

### Western blot

Cultured cells were lysed with ice-cold IP lysis buffer (Thermo Fisher Scientific, 87788) containing protease inhibitor cocktail (Sigma-Aldrich, 11836170001) and phosphatase inhibitor (Sigma-Aldrich, 04906845001) to prepare the total cell lysate. The protein concentration of the lysate samples was determined using a BCA assay (Thermo Fisher Scientific, 23227), and the input protein amount for western blot analysis was adjusted accordingly. The lysate samples were denatured by mixing with Blue Loading Gel Dye and DTT (Cell Signaling Technology, 7722). Primary antibodies used in this experiment were as follows: TSP-1, anti-Thrombospondin-1 clone A6.1 (Thermo Fisher Scientific, MA5-13398) at a 1:500 dilution, and GAPDH, anti-GAPDH clone 14C10 (Cell Signaling Technologies, #2118) at a 1:1000 dilution. Secondary antibody staining employed anti-mouse IgG and anti-rabbit IgG HRP-linked antibodies (Cell Signaling Technologies, #7076 and #7074, respectively) were used at 1:5000 dilution. The primary antibody staining step was performed overnight at 4°C, followed by the secondary antibody staining at RT for 1 hour. To visualize the protein size ladder, Precision Plus Protein™ WesternC™ Blotting Standards and Precision Protein™ StrepTactin-HRP Conjugate (Bio-Rad, 1610376 and 1610381) were used. Bullet Blocking One for Western Blotting (Nacalai USA, 13779-01) was used for blocking and as the antibody diluent. Western blot bands were visualized using Pierce™ ECL Western Blotting Substrate (Thermo Fisher Scientific, 32106) with the Odyssey FC imaging system (LI-COR Biotechnology), and the analysis was performed using Image Studio Software (v5.2.5, LI-COR).

### Bulk RNA-sequencing of *in vitro* cells

Total RNA was extracted from tumor cell pellets using RNeasy Mini Kit (Qiagen, 74106) and RNase-Free DNase Set (Qiagen, 79254) following the manufacturer’s protocol. RNA integrity was evaluated using Agilent Bioanalyzer 2100 and confirmed to be RIN 9.8 or greater. The following library preparation and sequencing were performed by DNA Technologies and Expression Analysis Core Laboratory at the University of California, Davis (UC Davis) Genome Center. Strand-specific and barcode-indexed RNA-Seq libraries were generated from 300 ng total RNA each after poly-A enrichment using the mRNA-Seq Hyper Kit (Kapa Biosystems, KK8581) following the manufacturer’s instructions. The fragment size distribution of the libraries was verified via microcapillary gel electrophoresis on a Bioanalyzer 2100. The libraries were quantified by fluorometry on a Qubit fluorometer (Life Technologies) and pooled in equimolar ratios. The pool was quantified by quantitative PCR with a Library Quant Kit (Kapa Biosystems, KK4824) and sequenced on an Illumina NovaSeq 6000 with paired-end 150 bp reads.

### Bulk RNA-sequencing of *in vivo* tumors

To perform RNA-seq with *in vivo* tumor tissue samples, mice were euthanized before being perfused via transcardial injection of 10 mL cold PBS. The whole brains were then harvested, and visible tumor tissues were dissected and stored in RNAlater (Invitrogen, AM7024) at 4°C until RNA extraction (within 1 week). Total RNA was extracted using the RNeasy Mini Kit (Qiagen, 74106) and RNase-Free DNase Set (Qiagen, 79254) following the manufacturer’s protocol. The tissues were homogenized using a Bioruptor standard water bath sonicator (Diagenode, UCD-200) with the following settings: power, H; duration, 5 min; cycle, 30s/30s. Due to the sonication treatment, the RIN scores were found to be low (4.2–6.5) when evaluated with an Agilent Bioanalyzer 2100. Because of these low RIN scores, we chose ribosomal depletion for mRNA enrichment. The subsequent library preparation and sequencing were performed by the UCSF Genomics CoLab. Starting material of 500 ng of total RNA was used according to vendor instructions with Universal plus mRNA with Nu Quant (TECAN, 0520), with noted changes to the protocol as follows. First, the Poly(A) Selection step was omitted. Second, we started with RNA fragmentation and used QIAseq FastSelect −rRNA/Globin Kit (Qiagen, 335377) to remove rRNA as follows: 0.1 µL of FastSelect rRNA and 1 µL of 1X Fragmentation Buffer was added to 10 µL of total RNA, with the addition of 10 µL of 2X Fragmentation Buffer. PCR program was used as follows: 94 °C/3min, 75 °C/2min, 70 °C/2min, 65 °C/2min, 60 °C/2min, 55 °C/2min, 37 °C/5min, 25 °C/5min, 10 °C/hold. After the rRNA depletion step, Universal plus mRNA with Nu Quant protocol was followed, starting with the first strand cDNA synthesis step but omitting the steps of AnyDeplete and NuQuant. After final library PCR amplification of 15 cycles and bead clean-up, individual libraries were pooled equally by volume, and quantified on Fragment Analyzer (Agilent, DNF-474). The quantified library pool was diluted to 1 nM and sequenced on MiniSeq (Illumina, FC-420-1001) to check for the quality of reads. Finally, individual libraries were normalized according to MiniSeq output reads, specifically by % protein-coding genes, and were sequenced on one lane of NovaSeq6000 SP PE150 (Illumina, 20028400) at UCSF Center for Advanced Technology (CAT).

### Bulk RNA-sequencing data processing and analysis

Quality checking, trimming, and removal of barcodes, were performed using fastp (version 0.20.0) with default parameters. The fastq reads were aligned to the mouse reference genome mm10 (GRCm38.p6) using STAR (v2.7.9a), with transcriptome annotation guidance from gencode.vM25.annotation.gtf. Sorting and indexing were performed using samtools (v1.14), and gene-level expression counts were estimated using stringtie (v2.0)^57^. The mapping status was visualized using the Integrative Genomic Viewer (IGV, version 2.8.0). All subsequent computational analyses were conducted using R (v4.1.2). Differential expression (DE) analyses comparing two groups were performed using the DESeq2 R package (v1.32.0) ^58^. The output was sorted based on the stat values to prepare the rank object. Preranked gene set enrichment analysis (GSEA) was performed using fgsea (1.18.0)^52^ for the following gene set collections: Hallmark, C2 KEGG, C2 Reactome, and C5 GO:BP (msigdbr 7.4.1).

### Mouse neonatal cortical neuron culture

Mouse neonatal cortical neuron cultures were prepared as described previously^8^. Neonatal (P1.5) C57BL/6J mice were euthanized by hypothermia followed by decapitation. The cerebral cortex was dissected under a microscope using aseptic techniques. Tissues from 4–6 mice were pooled and processed together. To isolate cortical neurons, the cerebral cortices were minced with scalpels on an ice-cold culture dish and digested using a papain dissociation kit (Worthington Biochemical, LK003150), according to the manufacturer’s instructions. Briefly, papain dissociation and subsequent protease inhibition were performed at 37 °C for 7 min and 3 min, respectively. The cell pellets were washed with Neurobasal-A medium (Gibco, 10888022) supplemented with deoxyribonuclease (1 mg/mL; Worthington, LS002007). The cell suspension was passed through a 40 µm filter, counted, and centrifuged. The culture media for maintaining neuronal cells, called neuron culture media (NCM), was prepared by mixing neurobasal-A medium with 1X B-27 (Gibco, 17504044), 1% Glutamax (Gibco, 35050061), 1% sodium pyruvate (Gibco, 11360070), and 1% Antibiotic-Antimycotic (Gibco, 15240062). The cell pellet was resuspended in neuron plating media (NPM), which was prepared by adding 4.5% fetal bovine serum (FBS) to the NCM. The cells were plated onto poly-D-lysine and laminin-coated coverslips (NeuVitro, GG-12-1.5-laminin) at a density of 1.5×10^5^ cells per well for 24-well plates (approximately 7.9×10^4^ cells per cm^2^). After 24 hours, the culture media was replaced with serum-free NCM. The cell culture was maintained by replenishing half of the culture media volume with fresh NCM every 3–4 days. For co-culture experiments involving neurons and SB28 glioblastoma cells for synapse imaging, cell suspensions of SB28-TSP-1-WT or KO cells (1.5×10^4^ cells) were added on top of the neurons maintained on the coverslips at day 7 of neuron culture, and maintained for 24 hours.

### Immunofluorescence

Immunofluorescence staining was performed according to standard methods. For *in vitro* samples, neurons and tumor cells were maintained on the poly-D-lysine-laminin-precoated coverslips (NeuVitro, GG-12-1.5-laminin or GG-18-1.5-Laminin) as mentioned above. For *in vivo* tissue samples, mice were euthanized before intracardiac perfusion with 10 mL cold PBS. The brains were harvested, fixed with 4% paraformaldehyde-PBS overnight, then soaked in 30% sucrose for one day or longer. Fixed brain tissues were subsequently embedded in Tissue-Tek® O.C.T. Compound (Sakura Finetek, 4583). Serial 10-mm coronal sections were then cut on freezing microtome and stored at −80°C until use. Cell samples or tissue sections were fixed with 10% formalin for 10 min, washed twice with 1× wash buffer (Dako, S300685-2C), blocked with PBS containing 5% normal goat serum (Abcam, ab7481) and 1% TruStain FcX PLUS anti-mouse CD16/32 Antibody (BioLegend, 156604) for 40 min at RT, then stained with primary antibodies overnight at 4°C. Primary antibodies used and the dilutions were as follows: for neurons: anti-Tubulin-βIII Rabbit mAb (1:200, BioLegend, 802001, clone TUJ1); anti-Synapsin-1 Mouse mAb (1:200, Synaptic Systems, 106 011); anti-Homer1 Guinea Pig pAb (1:200, Synaptic Systems, 160 004); for TSP-1 expressed on tumor-harboring mouse brains: anti-Thrombospondin-1 Rabbit pAb (1:200, ab85762, Abcam); After washing twice with 1X Dako wash buffer, staining with secondary antibodies was performed at the dilution of 1:250 for 1 hour at RT to detect primary labeling. The secondary antibodies used were as follows: for neurons: Goat anti-Rabbit IgG (H+L) AF514 (Invitrogen, A-31558); Goat anti-Mouse IgG AF594 (Abcam, ab150120); Goat anti-Guinea Pig IgG (H+L) AF647 (Invitrogen, A-21450); for TSP-1, Goat anti-Rabbit IgG H&L AF647 (Abcam, ab150083). After wash and dehydration, the stained samples were mounted with ProLong Diamond Antifade Mountant with DAPI (Fisher Scientific, P36971) on glass slides and covered with cover glasses.

### Tissue image acquisition and quantitative analysis

Images of *in vivo* tissue samples were acquired using a Zeiss Axio Imager 2 microscope (20× magnification) with TissueFAXS scanning software (TissueGnostics). Consistent exposure times and thresholds were maintained within each imaging session. Raw image data was imported into ImageJ software (version 2.9.0) for quantitative analysis. The tumor outline was manually identified in FITC channel-images for GFP signal detection and used as a region of interest (ROI). Signal intensity in the Cy5 channel-images for AF647 signal detection was measured within the ROI. Background signal intensities were measured in a non-tumor area of the same tissue. TSP-1 expression in the ROI was quantified as the adjusted mean fluorescence intensity, calculated as the mean signal intensities in the ROI minus the mean signal intensities of the background.

### Confocal imaging and colocalization analysis of synapsin-1 and homer-1 staining

Confocal images were captured at a resolution of 1,024 ×1,024 using a ×63 oil-immersion objective on a Zeiss LSM780 confocal microscope equipped with Zeiss Zen imaging software. The microscope settings for the synapsin-1 AF594 and homer-1 AF647 channels were kept constant across all samples throughout the experiment. The acquired Z-stack images were imporeted into ImageJ software (version 2.9.0) for further analysis. A 2D images were constructed from the Z-stack data using the Maximum Intensity Projection algorithm. Threshold values for the channels were consistently adjusted across all samples to ensure accurate identification of events. Colocalization events were defined as the overlap or adjacency between synapsin-1 puncta and homer-1-positive puncta. The colocalized puncta on neurites or between neuronal processes and SB28 glioblastoma cells were quantified manually using the Multi-point tool of ImageJ.

### Microelectrode array assay

To record spontaneous extracellular neuronal activity, a microelectrode array (MEA) assay was performed, as previously reported^8^. Briefly, we used Axion 24-well CytoView MEA plates, each well of which contains a 4×4 16 channel electrode array with channels spaced 350 µm apart, and the Maestro Edge system with integrated heating system and temperature controller (Axion Biosystems). The data acquisition and analysis were performed using an Axion Integrated Studio (AxIS). The neuronal firing events, or action potentials (i.e., “spikes”), and simultaneous bursts at multiple MEA electrodes (i.e., “network synchrony”) were measured using Axion Integrated Studio (AxIS) Navigator 3.5.2 software (Axion Biosystems) with the Neural Real-Time module. Weighted mean firing rate (WMFR) were defined as the spike rate per well multiplied by the number of active electrodes in the associated well. After data normalization, raster plots illustrating spike histograms and network bursts were generated using the Neural Metric Tool software (Axion Biosystems). The isolated mouse neonatal cortical neurons were plated and maintained as previously reported, and recordings were made at 24 hours after co-culture as indicated in the figure legends.

### Flow cytometry

Brain-infiltrating leukocytes (BILs) were isolated by density-gradient centrifugation using Percoll (GE Healthcare Life Sciences, 17089101), as previously described^59,60^. After isolation, single-cell suspensions of BILs (0.5–1×10^6^ cells) were stained with Fixable Viability Stain 780 solution (BD Biosciences, 565388) to discriminate live and dead cells. Fc receptor blocking was performed using TruStain FcX PLUS anti-mouse CD16/32 Antibody (BioLegend, 156604) prior to staining with fluorophore-conjugated antibodies at the concentrations recommended by the manufacturers. Fluorescence minus one (FMO) controls were used to determine the accurate gating. The antibody panels are listed in **Supplementary Tables S1–2**. Data was acquired using an Attune NxT flow cytometer (Thermo Fisher Scientific), and analyzed using FlowJo software (Tree Star, version 10.8.1).

**Supplementary Table 1.** Flow cytometry antibody panel for myeloid cells.

| Marker | Clone | Fluorophore | Manufacturer | Catalog ID | $\mu\text{L}$ for $5 \times 10^5$ cells |
| --- | --- | --- | --- | --- | --- |
| CD45 | 30-F11 | AF700 | BioLegend | 103128 | 0.25 |
| CD11b | M1/70 | APC/Cy7 | BioLegend | 101226 | 1 |
| F4/80 | BM8 | PE | BioLegend | 123110 | 0.5 |
| CD86 | GL-1 | PerCP-Cy5.5 | BioLegend | 105027 | 2.5 |
| CD206 | C068C2 | PE Dazzle | BioLegend | 141731 | 1 |

**Supplementary Table 2.** Flow cytometry antibody panel for T-cells.

| Marker | Clone | Fluorophore | Manufacturer | Catalog ID | $\mu\text{L}$ for $5 \times 10^5$ cells |
| --- | --- | --- | --- | --- | --- |
| CD45 | 30-F11 | AF700 | BioLegend | 103128 | 0.25 |
| CD3 | 17A2 | AF594 | BioLegend | 100240 | 0.5 |
| CD4 | RM4-5 | BV650 | BioLegend | 100546 | 2 |
| CD8 | 53-6.7 | BV605 | BioLegend | 100744 | 1 |
| CD25 | 3C7 | APC | BioLegend | 101909 | 0.5 |
| PD1 | 29F.1A12 | BV421 | BioLegend | 135221 | 0.5 |

## Statistical analysis

Mouse survival analysis was performed using the log-rank test from the R package survival, and Kaplan-Meier survival curves were visualized using the R package *survminer*. A significance level of *P* < 0.05 was regarded as statistically significant. For two-group comparisons, Wilcoxon rank-sum test or Student’s t-test was used, unless otherwise specified. For multiple testing, *P* values were adjusted using Benjamini-Hochberg or Bonferroni procedures, as specified in figure legends, and presented as adjusted *P* values. Statistical analysis and data visualization were performed with R version 4.1.2.

## Data availability

Transcriptome datasets generated during this study will be made available through the NCBI Gene Expression Omnibus (GEO) website. All other study data are included in the manuscript and/or supporting information.

## Acknowledgements

We thank the following individuals and organizations: the study participants and their families; Anny Shai and Yunita Lim of the UCSF Brain Tumor Center SPORE Biorepository and Pathology Core for their services (NIH/NCI 5P50CA097257-18); Mylinh Bernardi and Horng-Ru Lin of the Gladstone Genomics Core for their assistance with Visium FFPE Spatial Gene Expression assay and sequencing under the support from the James B. Pendleton Charitable Trust for funding NextSeq 500 sequencer used in this study; Braise Ndjamen of the Gladstone Histology and Light Microscopy Core for 10X Visium Spatial Transcriptomics data acquisition-related services; Anna Celli of the Laboratory for Cell Analysis Core Facility, UCSF Helen Diller Family Comprehensive Cancer Center for her technical assistance related to tissue image acquisition; Lenka Maliskova, Armita Norouzi, and Walter Eckalbar of UCSF Genomics CoLab for the RNA-sequencing library preparation; Sequencing was performed at the UCSF Center for Advanced Technology (CAT) for sequencing, supported by UCSF PBBR, RRP IMIA, and NIH 1S10OD028511-01 grants; the DNA Technologies and Expression Analysis Core at UC Davis Genome Center for bulk RNA-Seq services; members of the S.H.-J. laboratory and H.Okada laboratory for assistance, advice, and helpful discussions.

## Author contributions

Conceptualization: T.N., S.K., S.L.H.-J., and H.Okada; Formal analysis: T.N., and H.Okada; Funding acquisition: T.N., S.H.-J., and H.Okada; Investigation: T.N., S.K., C.J., A.Y., J.S.Y., S.L., T.C., S.S.S.P, D.D., A.G.S.D.; Methodology: T.N., S.K., A.Y., S.L., H.Ogino, P.W., S.L.H.-J., and H.Okada; Project administration: T.N., S.K., S.L.H.-J., and H.Okada; Resources: S.K., A.G.S.D., A.C., D.R.R., and S.L.H.-J.; Supervision: S.L.H.-J., and H.Okada; Visualization: T.N.; Writing (original draft): T.N., S.L.H.-J., and H.Okada; Writing (review and editing): T.N., S.K., A.Y., J.S.Y., P.W., D.R.R., S.L.H.-J., and H.Okada

## Prior publications

Part of this work was presented at the 2022 Annual Meeting of the Congress of Neurological Surgeons (CNS, 10/11/2022, San Francisco, CA, USA), at the 27th Annual Meeting and Education Day of the Society for Neuro-Oncology (SNO, 11/18/2022, Tampa, FL, USA), at the Annual Meeting of the American Association for Cancer Research (AACR, 04/17/2023, Orlando, FL, USA), at the 75th Annual Meeting of the American Academy of Neurology (AAN, 04/24/2023, Boston, MA, USA), and at the 20th Annual Meeting of the Association for Cancer Immunotherapy (CIMT, 05/06/2023, Mainz, Germany)

## Materials and Correspondence

All request for materials and correspondence may be sent to

## Extended Data

**Extended Data Fig. S1.**
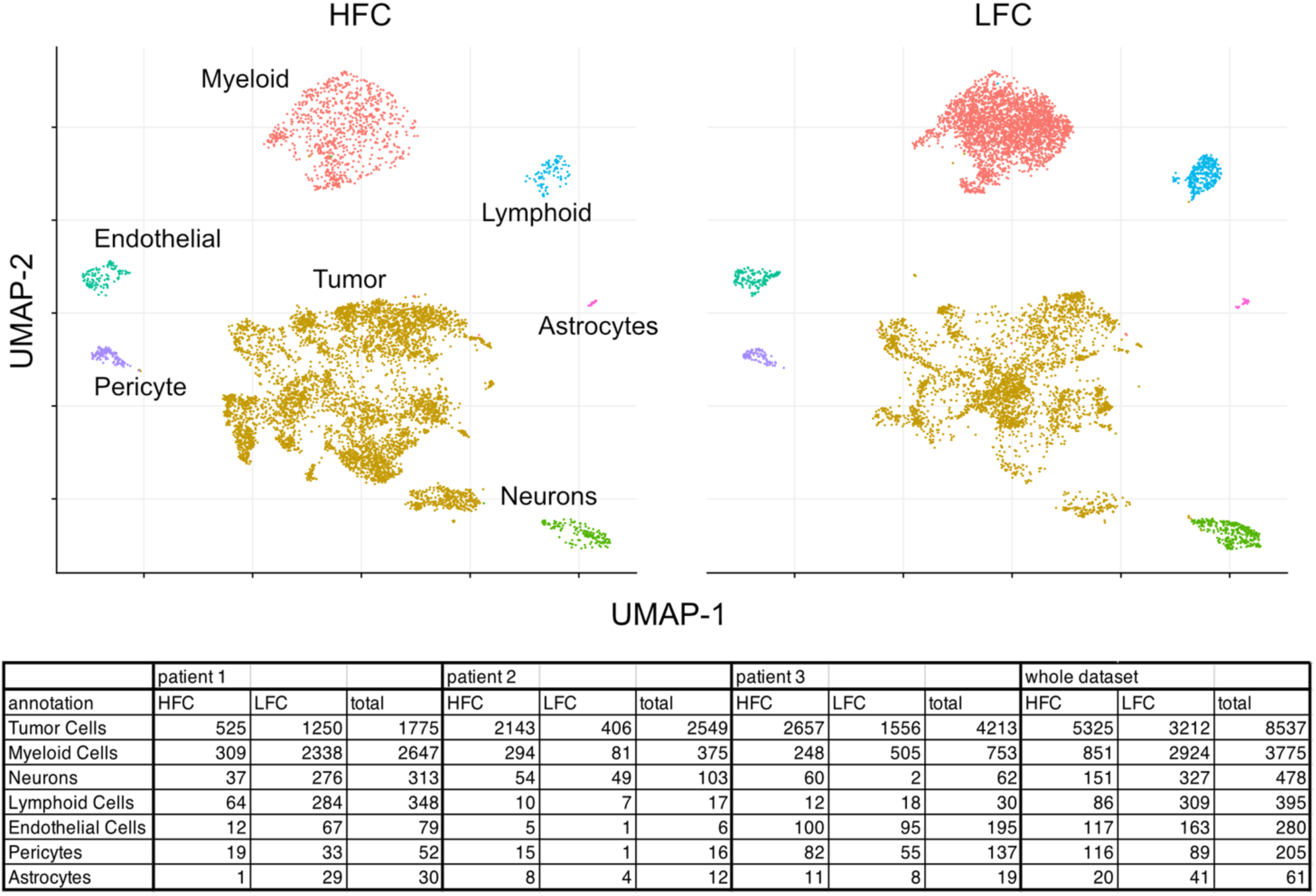
Overview of single-cell RNA-seq dataset of clinical samples annotated as either HFC- or LFC-derived (Related to Fig. 1) The UMAP plots showing each identified cluster, as previously reported^8^. The corresponding table shows the breakdown of the number of cells within the dataset.

**Extended Data Fig. S2.**
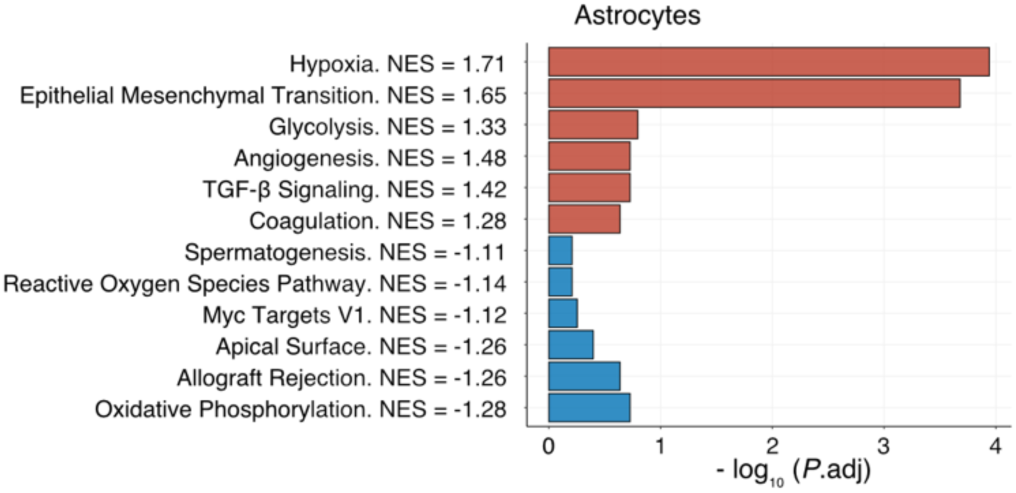
GSEA comparing HFC vs. LFC within astrocytes. (Related to Fig. 1) The top 6 upregulated and down-regulated signatures are shown. Red and blue indicate upregulation in HFC and LFC, respectively.

**Extended Data Fig. S3.**
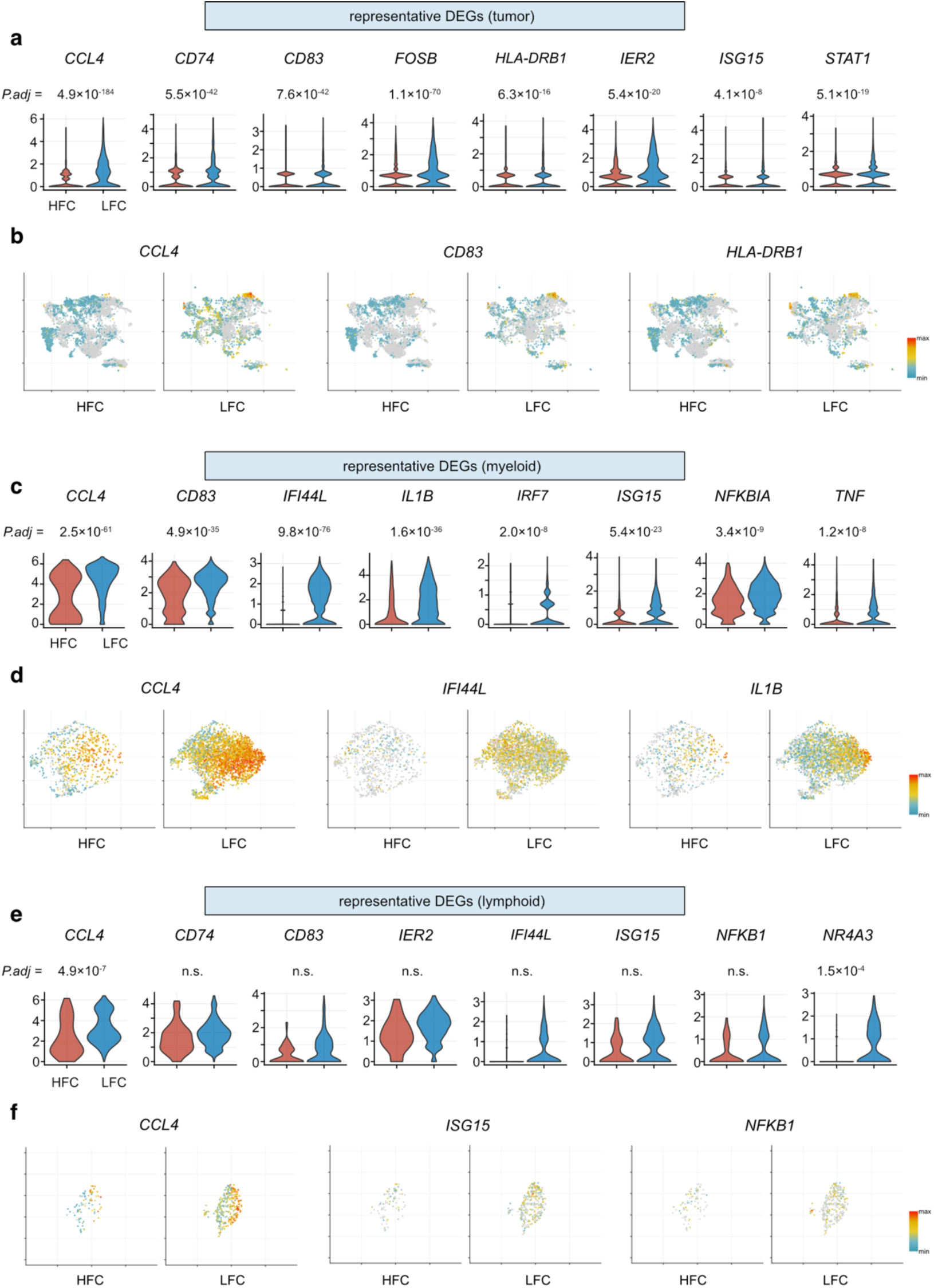
Representative differentially expressed genes (DEGs) in tumor, myeloid, lymphoid cells between HFC and LFC. (Related to Fig. 1) **a–f,** Violin plots (**a, c, e**), and feature plots (**b, d, f**) highlighting representative leading-edge genes differently expressed in tumor (**a–b**), myeloid (**c–d**), and lymphoid (**e–f**) cells between HFC and LFC regions. *P* values are calculated using the MAST algorithm with Benjamini-Hochberg adjustment.

**Extended Data Fig. S4.**
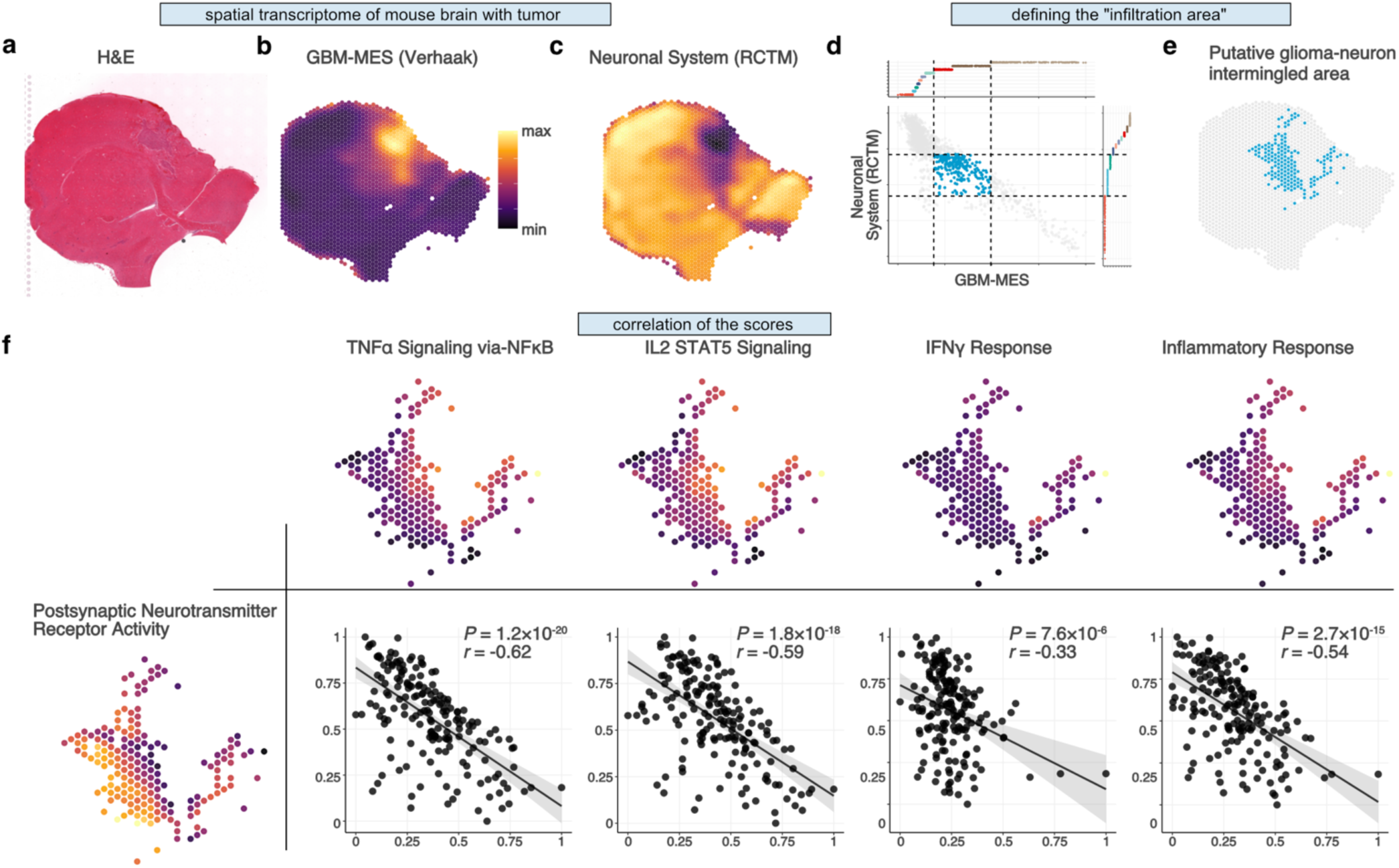
Spatial transcriptomic analyses of sample 2. (Related to Fig. 3) **a–c,** Histological images (H&E, upper) (**a**), and surface plots showing the distribution of the gene set enrichment scores of *Verhaak Glioblastoma Mesenchymal* (“GBM-MES” [C2:CGP]) (**b**) and *Neuronal System* (Reactome) (**c**). **d–e,** Scatter plot showing the relationship of the scores of *GBM-MES* and *Neuronal System* throughout the entire data set, where the spots with upper 10–30 percentiles of *GBM-MES* scores and lower 10–30 percentiles of *Neuronal Systems* (Reactome) scores are highlighted with blue (**d**). The surface plot shows the “putative glioma-neuronal infiltration areas” defined based on the distribution of the scores of *GBM-MES* and *Neuronal Systems* within the tumor bed (**e**). **f,** Surface plots show the gene set enrichment signature scores of *Post-synaptic Neurotransmitter Receptor Activity* (GO:MF), *TNFα-Signaling via NFκB*, *IL2-STAT5 Signaling*, *IFNγ Response*, and *Inflammatory Response* (all from Hallmark) within the defined putative glioma-neuronal intermingled area. Scatter plots show the correlations between the scores of *Post-synaptic Neurotransmitter Receptor Activity* and the others. *P* values are calculated using Pearson correlation test. *r*, correlation coefficient.

**Extended Data Fig. S5.**
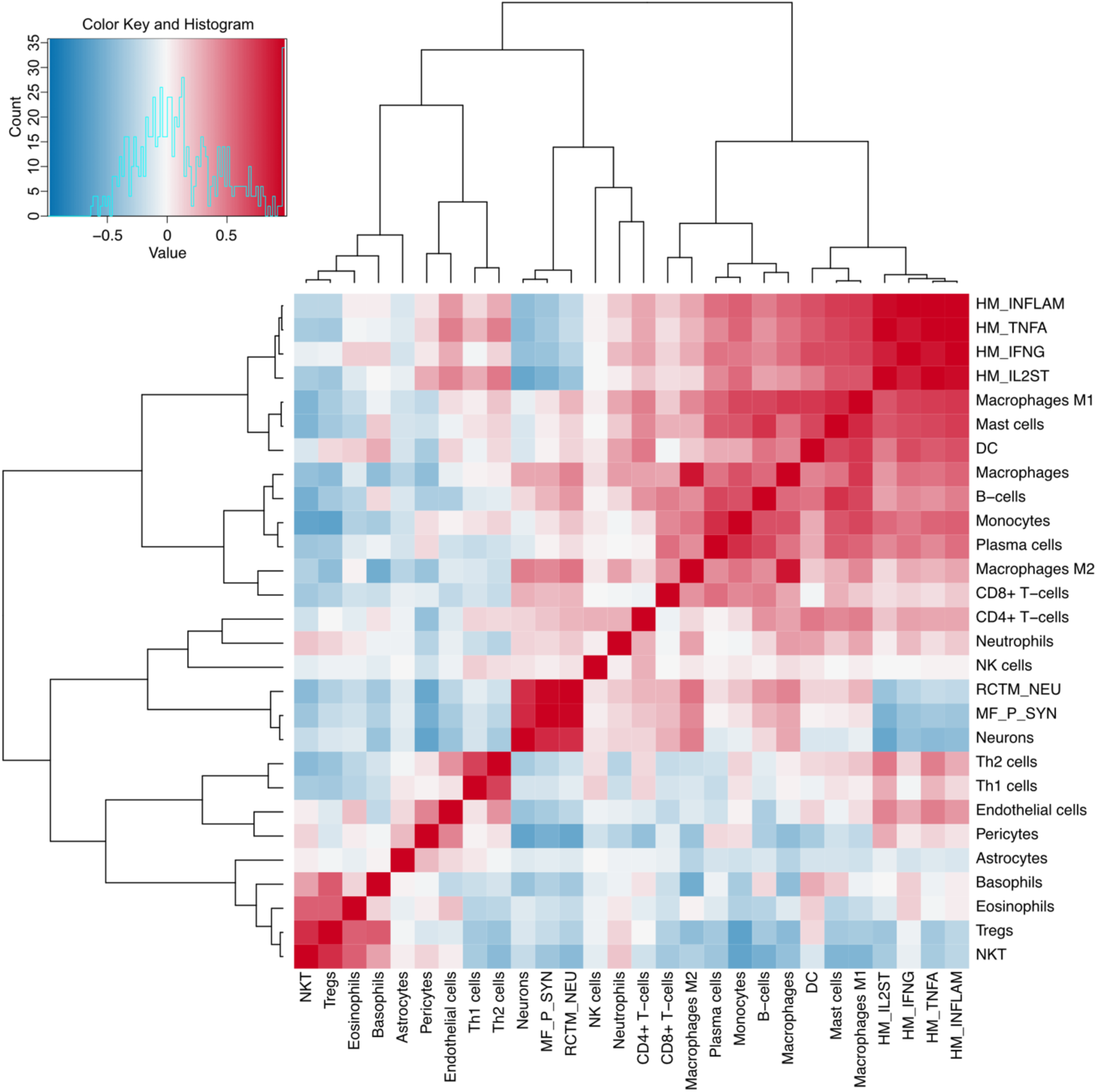
Hierarchical clustering heatmap of pairwise correlations among the estimated cell abundances and gene sets. (Related to Fig. 3) Cell-type deconvolution analysis is performed on the transcriptome data of each spot within the glioma-neuron infiltration area (n = 148 spots) using xCell^56^. Pearson’s correlations are evaluated among the estimated scores of the curated 22 cell types from xCell and additional 6 gene signature scores. Cell types and signature scores are reordered according to hierarchical clustering. In the heatmap, blue and red indicate positive and negative correlations, respectively.

**Extended Data Fig. S6.**
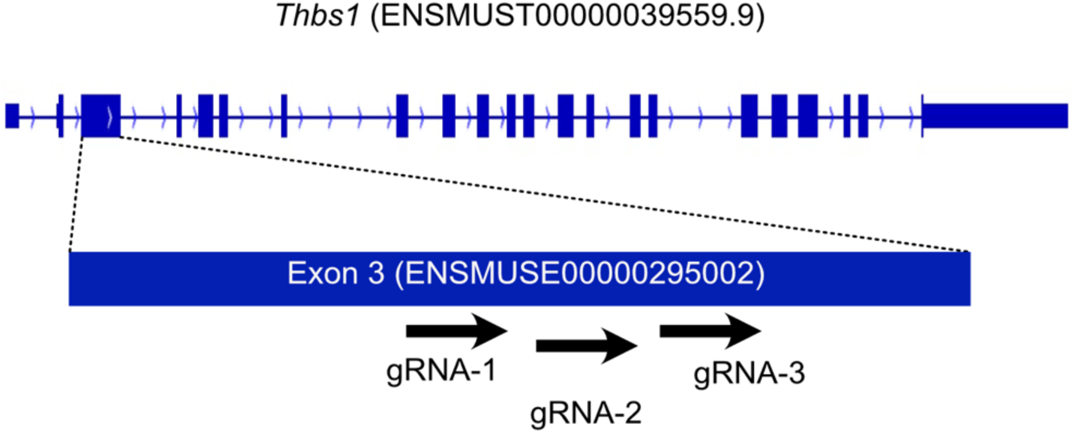
Design of CRISPR-KO targeting *Thbs1* gene. (Related to Fig. 4) A scheme showing the design of the Gene KO kit v2 (Synthego) containing three multi-guide sgRNAs specifically targeting regions within exon 3 (ENSMUSE00000295002) of the murine *Thbs1* gene.

**Extended Data Fig. S7.**
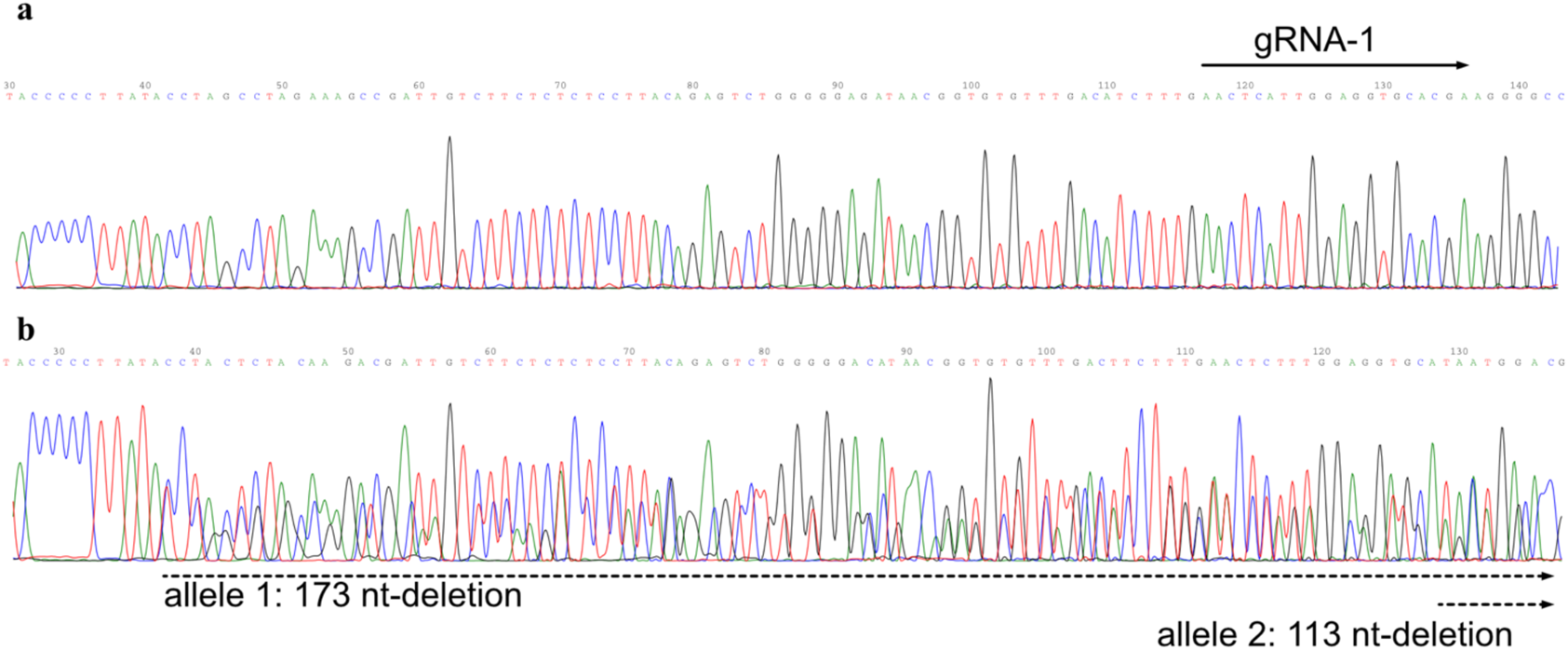
Sanger sequencing of genomic DNA testing the *Thbs1* gene status. (Related to Fig. 4) Snapshot of Sanger sequencing data of genomic DNA extracted from SB28-TSP-1-WT (**a**) and KO (clone 1C1) (**b**).

**Extended Data Fig. S8.**
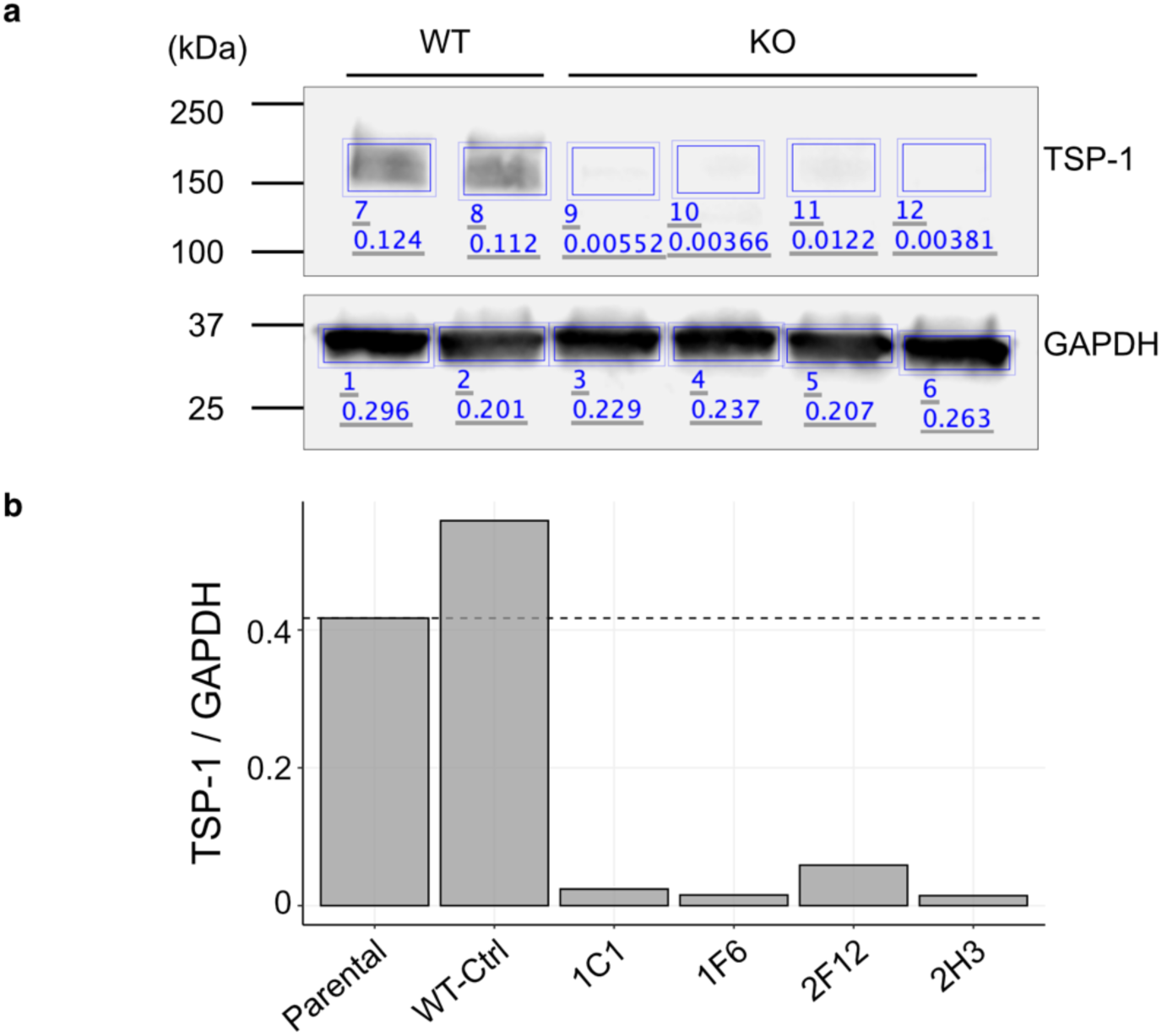
Western blot analysis of tumor cell lysate samples measuring the TSP-1 protein expression. (Related to Fig. 4) **a,** A western blot image showing each ROI on which signal intensity was measured. **b,** A bar plot showing the signal ratio of TSP-1 to GAPDH. Signal values were calculated as sum of the pixel intensity values (Total) for a shape minus the product of the Background and the Area (Signal = Total - [Background × Area]).

**Extended Data Fig. S9.**
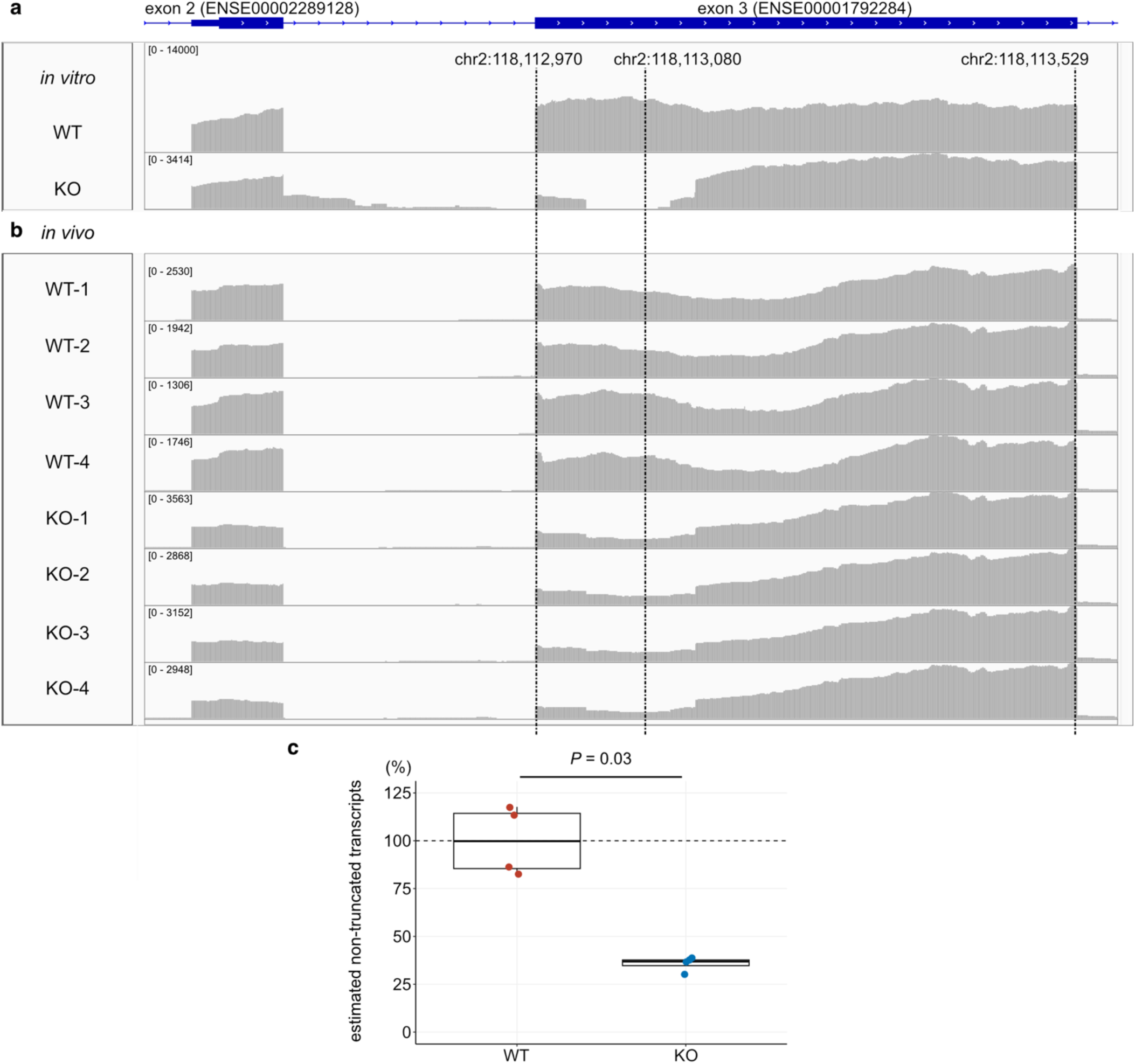
IGV snapshots showing RNA-seq reads mapped on the Thbs1 gene exon 3 region. (Related to Fig. 4) **a–b,** Snapshots of integrative genome viewer (IGV) showing the regions of exons 2–3 of the *Thbs1* genes and the mapped RNA-seq reads of *in vitro* culture cells (**a**) and *in vivo* mouse brain tumors SB28-TSP-1-WT and KO (n = 4 per group) (**b**). Break lines indicate the positions of the start of the exon (chr2:118,112,970), the middle of the deleted region (chr2:118,113,080), and the end of the exon (chr2:118,113,529). **c,** Box plot showing the estimated percentage of non-truncated (canonical) transcripts in each *in vivo* sample calculated as the read depth at the middle of the deletion divided by the average read depth at the start and the end of the exon and adjusted for the average of the WT cases to be 1. In the KO group, 70.9% of RNA-seq reads mapped on the *Thbs1* exon 3 region are estimated to be truncated (range, 68.1%–77.4%). *P* value is calculated using Wilcoxon rank-sum test.

**Extended Data Fig. S10.**
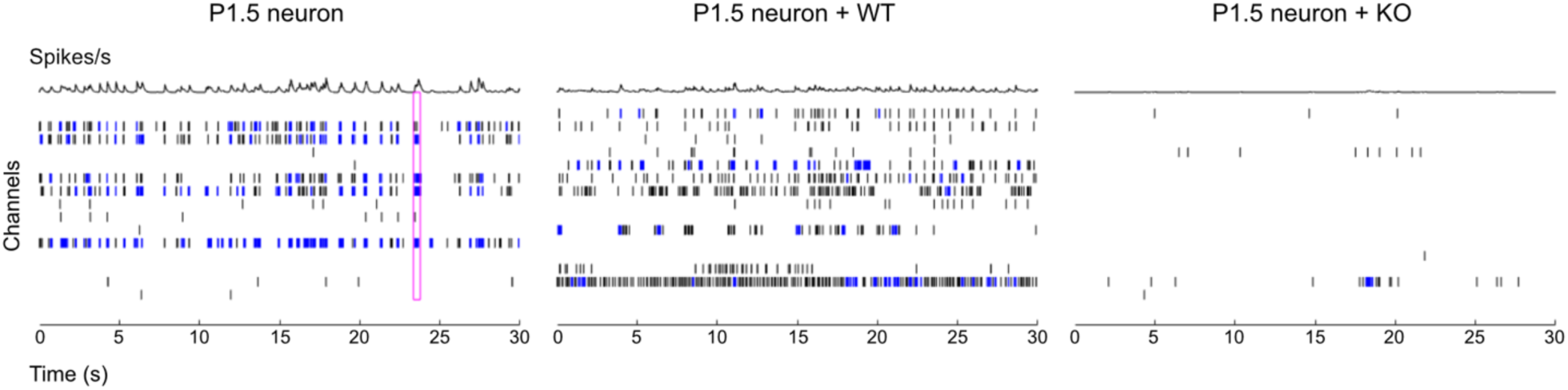
Representative MEA raster plots of mouse neonatal cortical neuron culture. (Related to Fig. 4) Representative MEA raster plots of mouse neonatal cortical neurons (left), neurons co-cultured with SB28-TSP-1-WT (middle) or KO tumor cells (right), recorded after 24 h co-culture. Each color indicates neuronal spikes (black tick marks), bursts (cluster of spikes in blue), and synchronized network bursts (pink).

**Extended Data Fig. S11.**
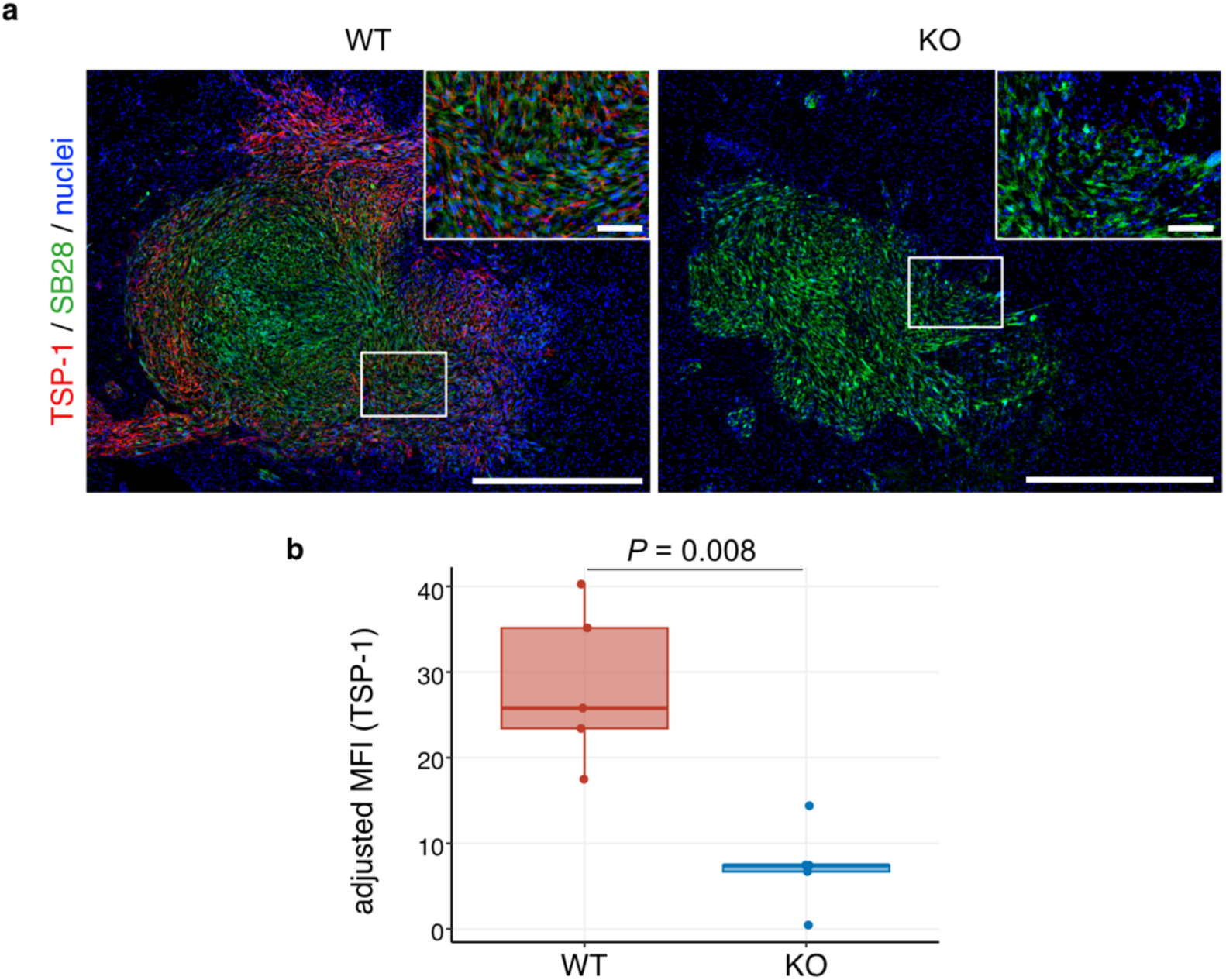
Representative immunofluorescence images of *in vivo* tumor lesions of SB28-TSP-1-WT or KO. (Related to Fig. 4) **a,** Representative immunofluorescence images of tumor lesions of SB28-TSP-1-WT (left) or KO (right) growing in syngeneic mouse brains. Red, TSP-1; green, GFP (SB28 tumor cells). Scale bar, 200 and 20 µm in low- and high-magnification images, respectively. **b,** Box plot showing the adjusted mean fluorescence intensity of TSP-1 signals measured in the GFP-positive tumor area adjusted by the background on the same tissue images. *P* value is calculated using Wilcoxon rank-sum test.

**Extended Data Fig. S12.**
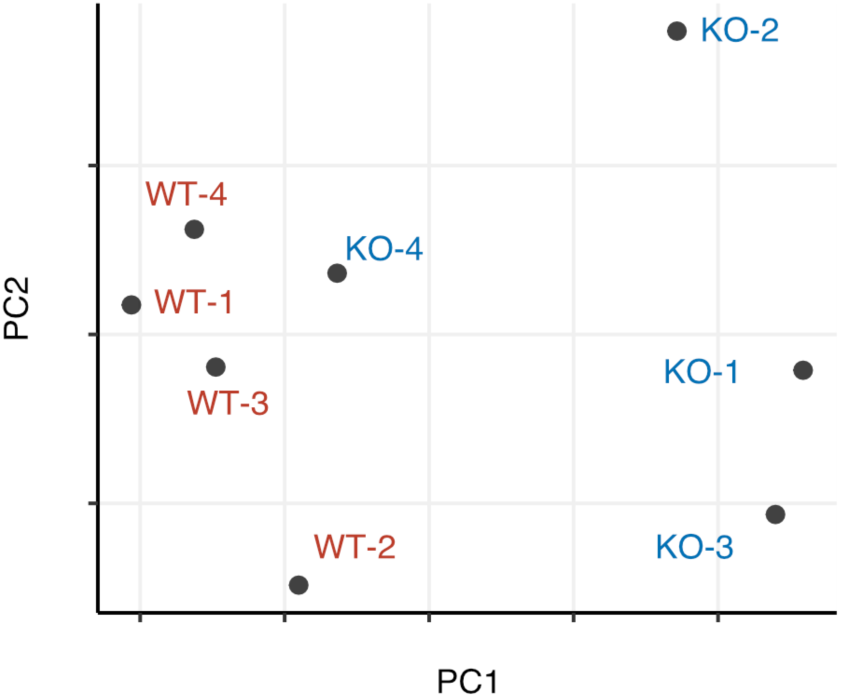
PCA plot showing the distinct gene expression patterns between in vivo tumors of SB28-TSP-1-WT and KO. (Related to Fig. 4) Principal component analysis (PCA) plot showing the similarities and differences in transcriptome of i*n vivo* tumor samples of SB28-TSP-1-WT (red) and KO (blue) groups (n = 4 mice per group).

**Extended Data Fig. S13.**
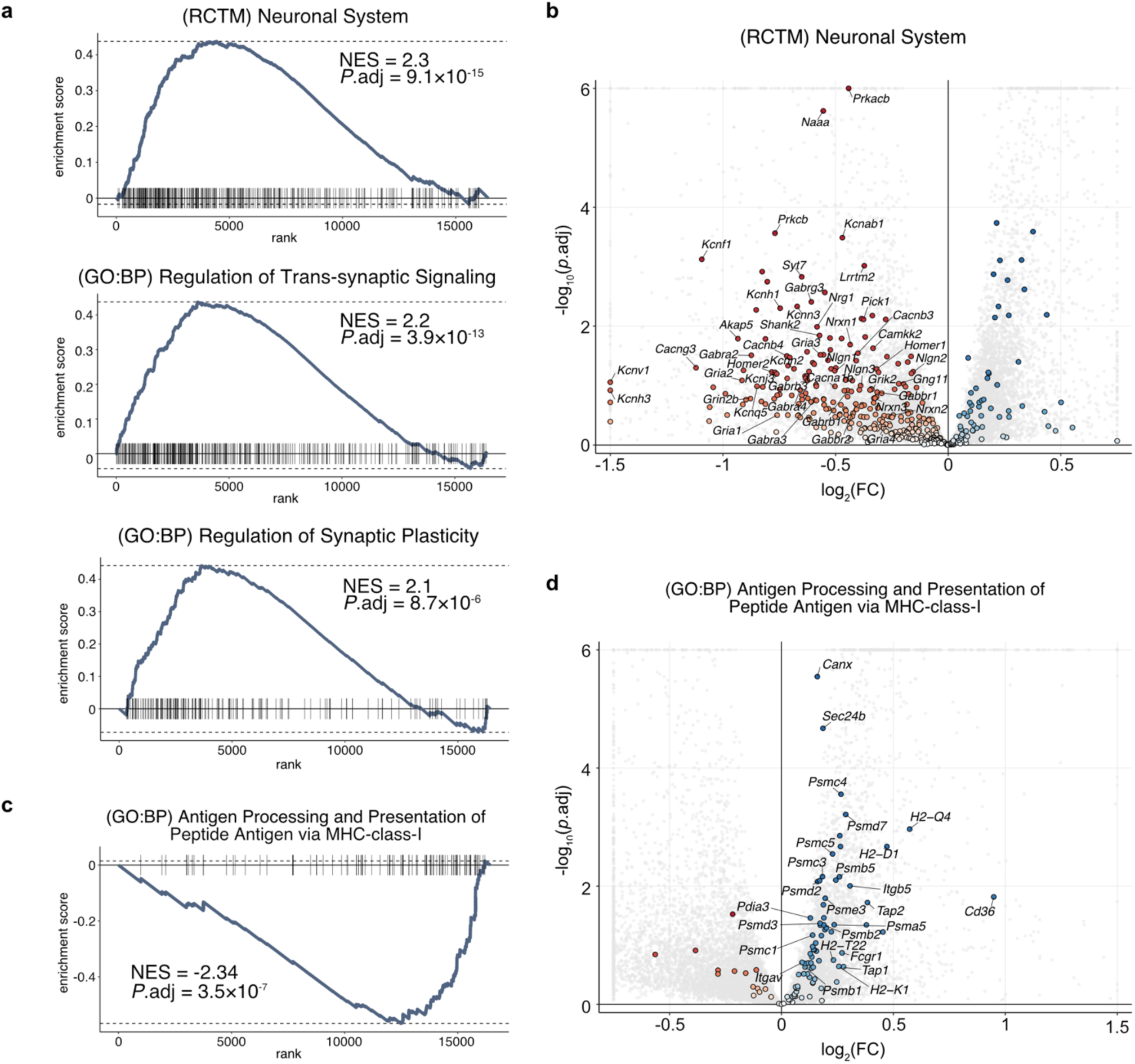
Representative pathways differentially expressed between *in vivo* SB28-TSP-1-WT and KO tumors. (Related to Fig. 4) **a and c,** Enrichment plots summarizing GSEA with the gene sets *Neuronal System* (Reactome), *Regulation of Trans-synaptic Signaling* (GO:BP), *Regulation of Synaptic Plasticity* (GO:BP) (**a**), and *Antigen Processing and Presentation of Peptide Antigen via MHC class I* (GO:BP) (**c**) to compare the gene expression patterns between the SB28-TSP-1-WT and KO mouse tumors. Positive and negative normalized enrichment scores (NES) indicate the upregulation and downregulation in WT tumors compared to the KO counterparts, respectively. **b and d**, Volcano plots showing the genes composing the gene set *Neuronal System* (Reactome) (**b**) and *Antigen Processing and Presentation of Peptide Antigen via MHC class I* (GO:BP) (**d**). The genes composing each gene set are highlighted in colors, and among them, the representative leading-edge gene symbols are labeled. Genes with log_2_FC values and adjusted *P* values exceeding the boundaries are flattened and shown on the edges.

**Extended Data Fig. S14.**
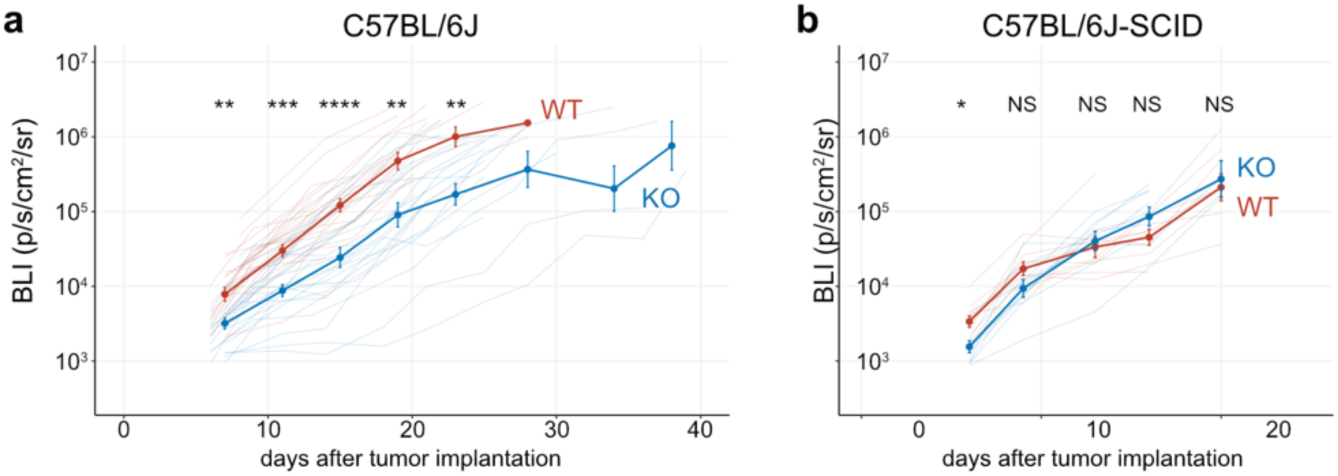
*In vivo* tumor growth curves. (Related to Fig. 4) **a–b,** Tumor size curves monitored by luciferase bioluminescence imaging (BLI) over time for the SB28-TSP-1-WT or KO inoculated into C57BL/6J immunocompetent mice (n = 25 mice each) (**a**), and immunocompromised C57BL/6J-SCID mice (n = 10 mice each) (**b**). Thin lines show traces for individual animals, and thick lines show geometric means. *P* values are calculated for each time point using t-test adjusted with Bonferroni’s multiple testing corrections. \**P* < 0.05, \*\**P* < 0.01, \*\*\**P* < 0.001, \*\*\*\**P* < 0.0001; NS, not significant.

**Extended Data Fig. S15.**
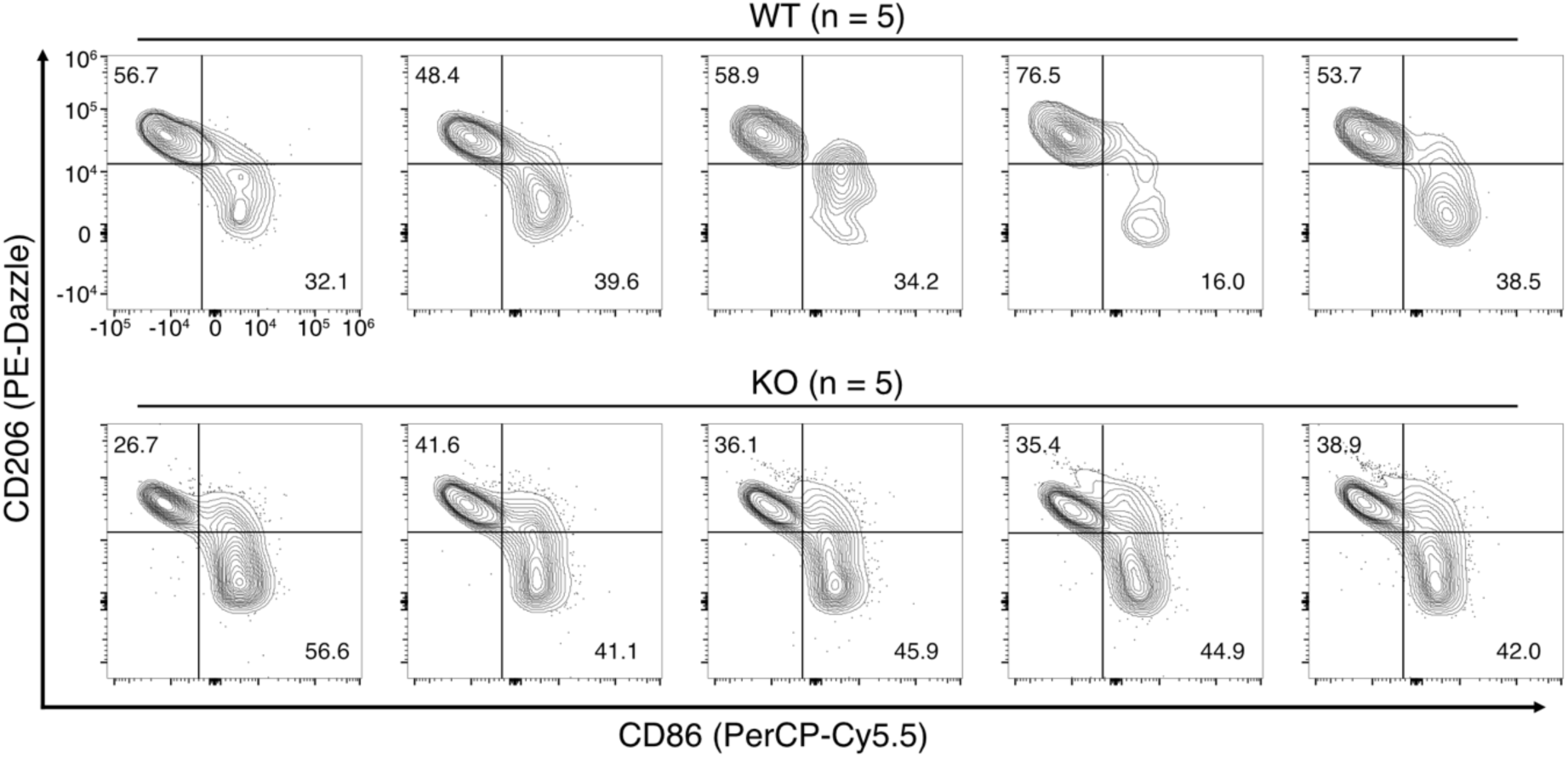
Distributions of classically activated and alternatively activated tumor-associated macrophage populations isolated from tumor-harboring mouse brains. (Related to Fig. 4) Contour plots showing the distributions of expression of CD86 and CD206 on the CD45+/CD11b+/F4-80+ TAMs isolated from each individual mouse brain harboring SB28-TSP-1-WT or KO cells (n = 5 mice per group). Values in the plots indicate the percentages of the cell populations identified within the gate.

**Extended Data Fig. S16.**
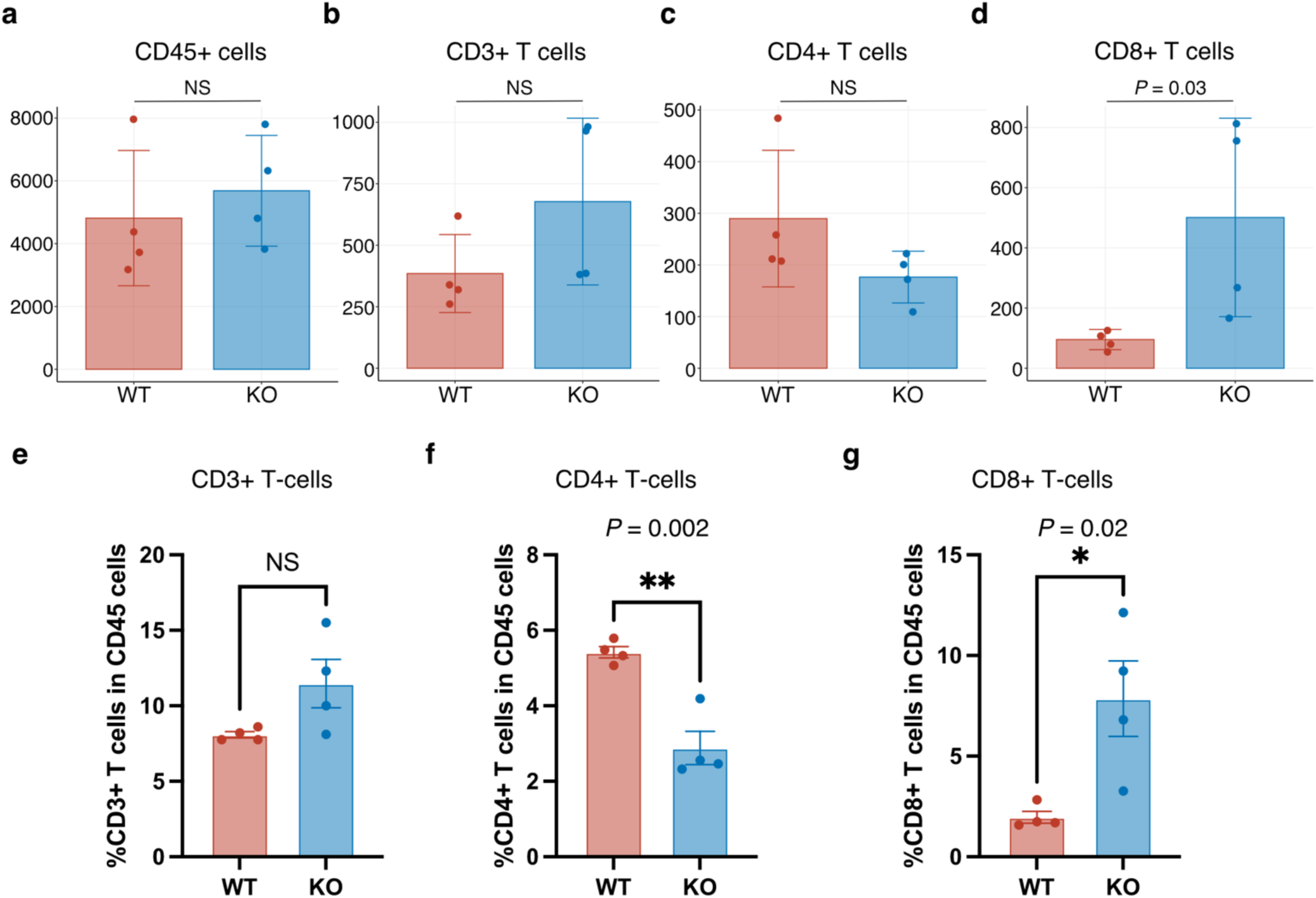
Abundances and fractions of T-cells in brain-infiltrating leukocytes. (Related to Fig. 4) **a–d,** Bar plots showing the abundance of CD45+ (**a**), CD45+/CD3+ (**b**), CD45+/CD3+/CD8+ (**b**), and CD45+/CD3+/CD4+ (**d**) T-cells identified in the samples. **e–g,** Bar plots showing the percentages of CD3+ (**e**), CD3+/CD8+ (**f**), and CD3+/CD4+ (**g**) T-cells within CD45+ cell populations.

**Extended Data Fig. S17.**
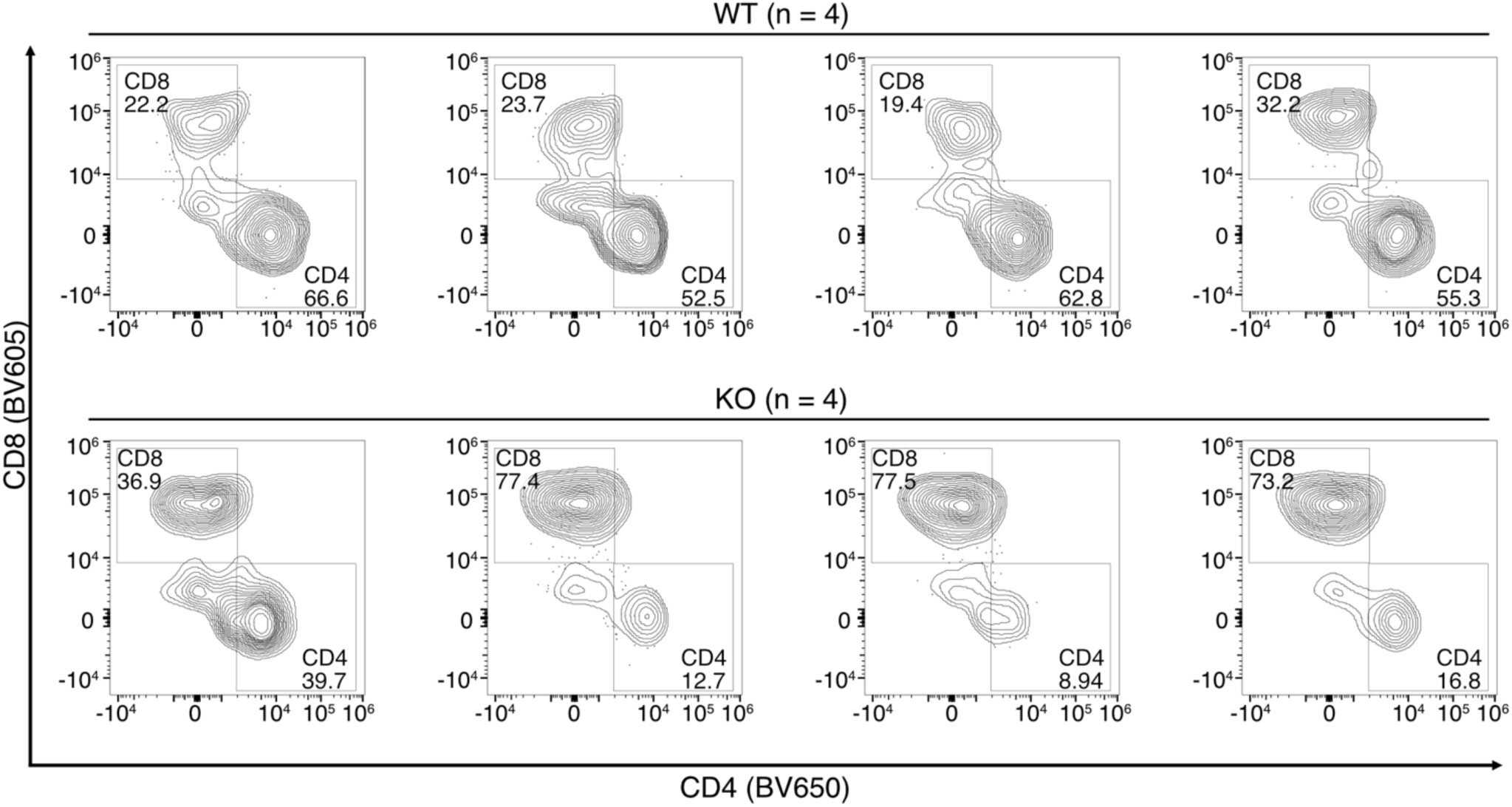
Distributions of brain-infiltrating CD4+ and CD8 + T-cell populations. (Related to Fig. 4) Contour plots showing the distributions of CD4+ and CD8+ cells within CD45+CD3+ cell populations in each individual sample (n = 4 mice per group). Values in the plots indicate the percentages of the cell populations identified within the gate.

**Extended Data Fig. S18.**
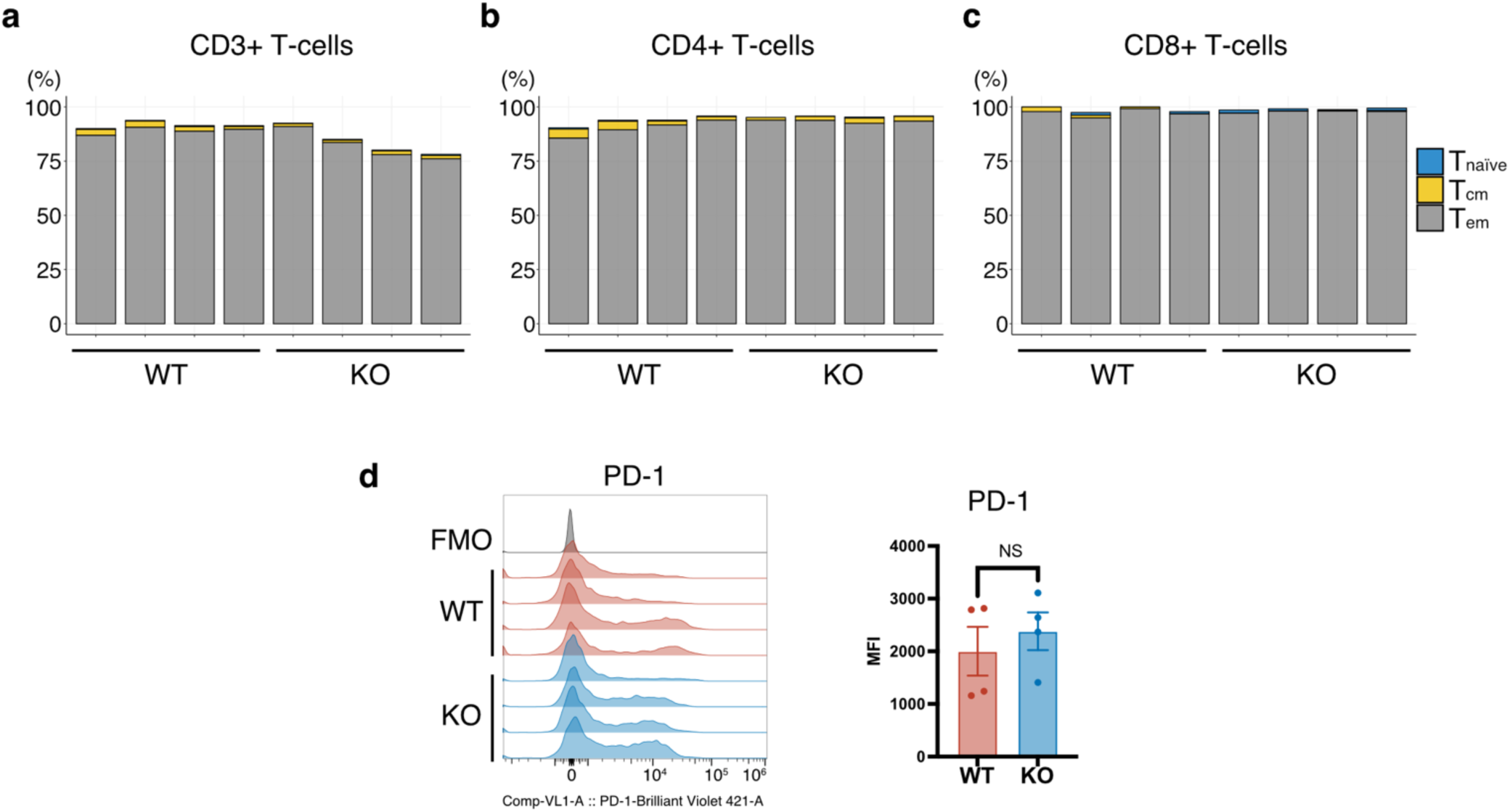
Characterization of brain-infiltrating T-cells. (Related to Fig. 4) **a–c,** Bar plots showing the fractions of naive (blue), central-memory (yellow), and effector-memory (gray) T-cell phenotypes detected in each sample.

**Extended Data Fig. S19.**
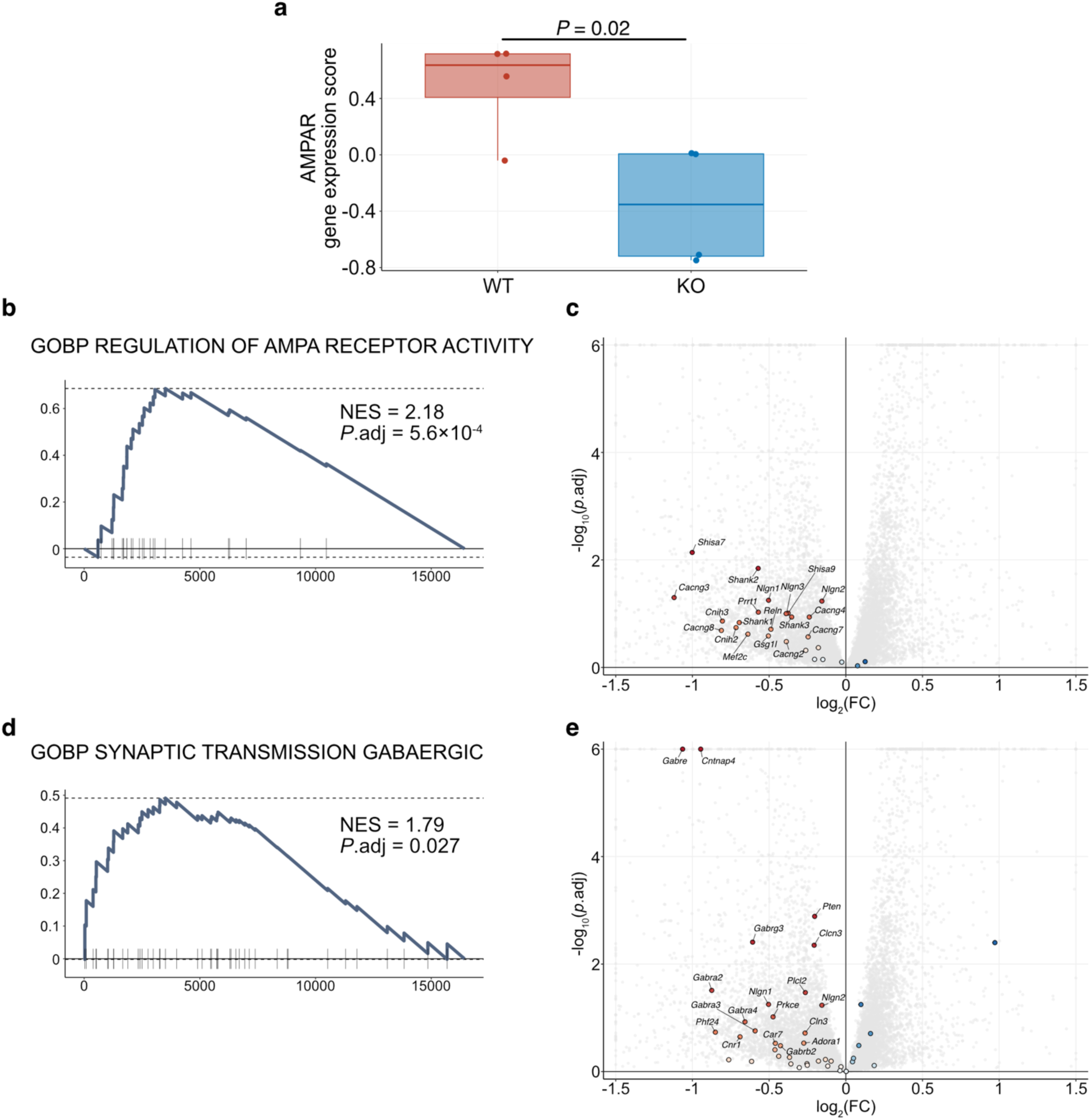
Pathway analyses of AMPAR and GABA-related signals. (Related to Fig. 4) **a,** Box plot showing the AMPA-R gene expression scores calculated based on bulk RNA-seq data of *in-vivo* tumors with TSP-1-WT and KO (n = 4 each). *P* value is calculated using t-test. **b–e,** Enrichment plots summarizing GSEA with the gene sets *Regulation of AMPA Receptor Activity* (GO:BP) (**b**), *Synaptic Transmission Gabaergic* (GO:BP) (**d**) to compare the gene expression patterns between the SB28-TSP-1-WT and KO tumors. In the corresponding volcano plots (**c**, and **e**), genes composing the gene sets are highlighted in colors, and the identified representative leading-edge genes are labeled with the gene symbol names. Background gray dots are all the protein-coding genes in the dataset. The genes composing each gene set are highlighted in colors, and among them, the representative leading-edge gene symbols are labeled. Genes with log_2_FC values and adjusted *P* values exceeding the boundaries are flattened and shown on the edges. NES, normalized enrichment score; FC, fold change **d,** Left: Histogram showing the distribution of fluorescence intensity of PD-1 measured in each sample (n = 4, each group). Right: Bar plot summarizing the median fluorescence intensity (MFI). FMO, fluorescence minus one control.

**Extended Data Fig. S20.**
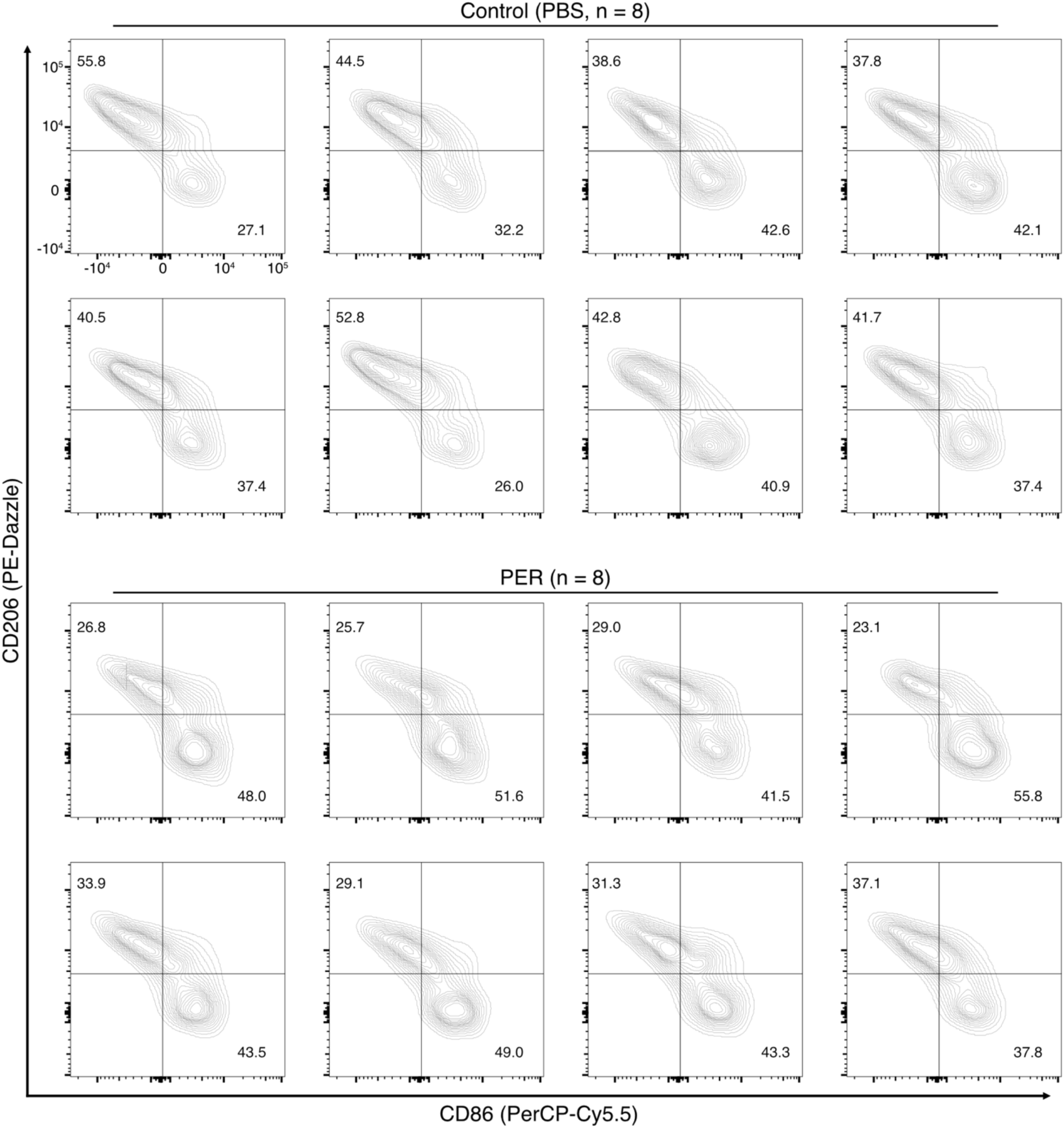
Distributions of classically activated and alternatively activated TAMs isolated from tumor-harboring mouse brains with and without PER treatment. (Related to Fig. 4) Contour plots showing the distributions of expression of CD86 and CD206 on the CD45+/CD11b+/F4-80+ TAMs isolated from brains of each individual SB28-TSP-1-WT harboring mouse treated with PER or control (PBS) (n = 8 mice per group).

